# Shared and Distinct High-Dimensional Object Spaces in Human and Macaque Inferotemporal Cortex

**DOI:** 10.64898/2026.05.20.724014

**Authors:** Sander van Bree, Martin N. Hebart

## Abstract

Human and macaque studies of inferotemporal cortex (IT) have shaped our understanding of object vision, yet the extent of their representational alignment and the precise nature of this correspondence have remained unclear. To address these questions, we applied multi-variate cross-species alignment to large-scale macaque multi-unit recordings and human fMRI responses to the same 8,640 naturalistic object images. The resulting shared space revealed a surprisingly rich, high-dimensional geometry that extended well beyond coarse correspondences such as animacy and shape and included a wide range of interpretable visual and conceptual dimensions, as well as specific conjunctions of the two. Unexpectedly, the space also uncovered a shared neural signature for orthography-related content, consistent with precursor mechanisms in macaques. Contrasting the within-species spaces revealed systematic differences in the expression of living and artificial categories, conspecific content, and visual versus conceptual structure. Together, these findings uncover a high-dimensional shared object space along with species-specific divergences, clarifying the scope and limits of cross-species comparability.

## Introduction

Humans and non-human primates make sense of the world by forging representations from an evolving stream of retinal inputs. Essential to this cascade is inferotemporal cortex (IT), a collection of regions situated in the ventral visual pathway. Extensive research has shown that neuronal responses in IT are sensitive to a range of features extracted from the visual inputs, including the shape (Tanaka, 1996) and identity of visual contents (Hung et al., 2005). By encoding shape, identity, and other properties such as color (Chang et al., 2017), IT exhibits a functional organization of representations that supports invariant object recognition (DiCarlo and Cox, 2007) and adaptation to the environment more broadly (Arcaro and Livingstone, 2021).

Despite extensive work in both humans and macaques (Arcaro and Livingstone, 2021; Conway, 2018; Mantini et al., 2012), we still lack a detailed understanding of the degree to which the functional organization of IT is shared across primate species. Prior work has identified coarse-grained correspondences, including sensitivity to visual properties such as shape and color (Denys et al., 2004; Sawamura et al., 2005; Lafer-Sousa et al., 2016), category-selective responses to faces and body parts (Bell et al., 2009; Pinsk et al., 2009; Tsao et al., 2008), dimensions related to animacy and shape (Kriegeskorte et al., 2008; Mur et al., 2013; Bao et al., 2020; Coggan and Tong, 2023; Yao et al., 2023), and shared representational structure in responses to natural scenes (Vinken et al., 2025; Li et al., 2026). However, these comparisons have typically examined predefined features, focused on category-selective regions, or sampled only a limited portion of object space. Consequently, beyond broad correspondences between humans and macaques, the rich shared organization of primate IT remains largely uncharacterized, including the degree to which this shared geometry is interpretable and to what extent its dimensions are visual or conceptual in nature. Furthermore, relatively little is known about how far these cross-species correspondences extend, and at which point differences in anatomy, experience, and ecological pressures give rise to divergent object representations. Addressing these gaps is important for understanding the comparability of representations between species and the inferences that can be drawn from scientific findings in either species alone.

To characterize the shared and species-specific organization of IT more comprehensively, here we developed a multivariate framework for separating shared from species-specific representational structure, drawing on densely sampled macaque multi-unit recordings and human fMRI responses from IT in response to 8,640 naturalistic object images spanning 720 categories. Using this framework, we identified a surprisingly rich and broadly interpretable representational geometry spanning primate IT that far exceeded established dimensions such as animacy and shape. Within this geometry, we identified a shared set of visual and conceptual dimensions, and dimensions reflecting specific combinations of the two. This reveals that in both species, there is similar visual and conceptual information that is intertwined—suggesting that IT organization binds the two forms of information. In addition, we identified shared structure related to orthographic stimuli, consistent with a shared precursor to the human visual representation of symbols. Comparisons of the within-species spaces further revealed systematic differences, specifically in the preferences for particular categories and conspecific content, as well as the relative balance between visual and conceptual information. Together, these findings reveal the high-dimensional organization of primate IT and delineate both its common organization across species and dimensions along which measured human and macaque object representations diverge.

## Results

### Defining shared and within-species IT spaces

To uncover the nature of primate IT representations and determine the degree of their generalization across species, we capitalized on two large-scale datasets collected in human and macaque subjects viewing the same broad sample of natural object space (Fig. 1). For the monkey data, we used the THINGS ventral stream spiking dataset, which measures multi-unit activity from V1, V4, and IT of two macaque monkeys (Papale et al., 2025). IT responses were averaged within a 75–175 ms post-stimulus window, which captured a temporally stable code (Fig. S1). For the human data, we used the THINGS-fMRI dataset, with data from three participants across 12 sessions each (Hebart et al., 2023; Contier et al., 2024). Together, this yielded five subject-specific datasets arranged as stimulus-by-feature matrices, where features correspond to IT electrodes for macaques and IT voxels for humans (Fig. S1; see Methods for ROI definition).

**Fig. 1.**
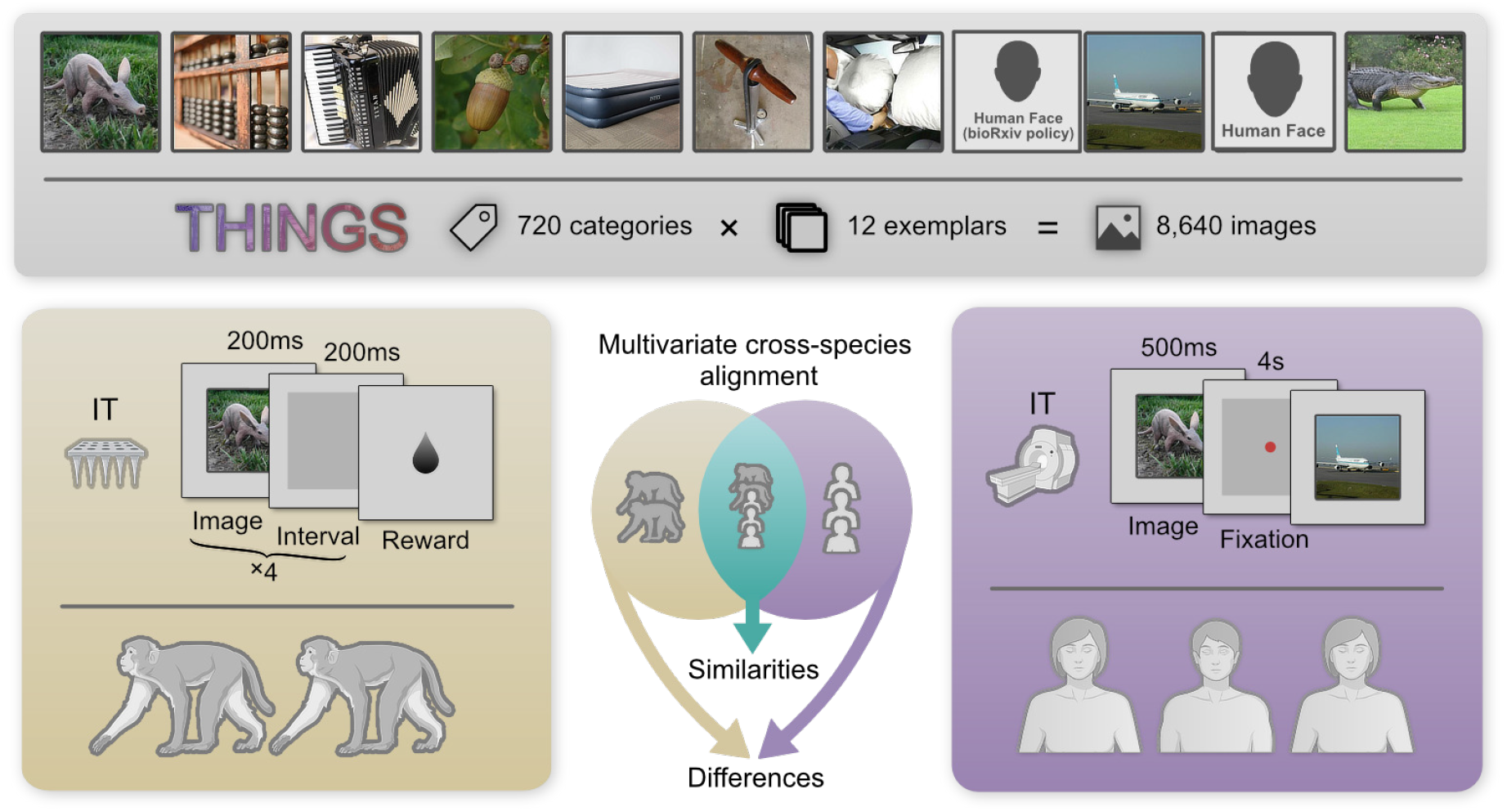
Stimuli, neuroimaging datasets, and main analysis framework. Macaques and human participants viewed naturalistic images from the THINGS image database while their brain responses were recorded. We compared multi-unit responses sampled from electrode arrays across monkey IT with fMRI responses from human IT. Neural responses to a shared set of 8,640 images (720 basic object categories with 12 exemplars) were aligned into three canonical spaces, targeting latent patterns shared across and within primate species.

In order to separate shared from species-specific representational structure, we developed an analysis framework for cross-species alignment. As a first step, we applied multiset canonical correlation analysis (MCCA), which takes advantage of the rich, multivariate structure present in both datasets. Specifically, MCCA estimates a set of linear transformations across multiple datasets that maximize cross-dataset correlations (Correa et al., 2010; Hardoon et al., 2004; de Cheveigné et al., 2019), highlighting shared structure that may be less apparent in analyses of individual subjects or in group-average models. We fit three models: a universal model across all five subjects, and separate within-species models for macaques (*n* = 2) and humans (*n* = 3). To discriminate representational structure from noise, we computed heldout cross-view correlations, retaining only those components that exceeded a permutation-derived null distribution.

This procedure uncovered 90 significant components for the cross-species alignment (FDR-adjusted *p <* 0.05; Fig. 2A). This number substantially exceeds prior characterizations that emphasized a core set of categories (Kriegeskorte et al., 2008) or a low-dimensional object space (Bao et al., 2020; Coggan and Tong, 2023), showing that, instead, primate IT carries a broad geometry composed of many reliable dimensions. Analyzing the within-species models supports this result by showing that the alignment across species yielded a higher-dimensional space relative to the alignment between humans (within-human components: 32; within-macaque components: 91, all FDR-adjusted *p <* 0.05; Fig. 2A). Together, these results highlight how between-species alignment can reveal representational structure that cannot be recovered in humans alone. In the following, we refer to the resulting spaces as canonical spaces, which formed the basis for all downstream analyses.

**Fig. 2.**
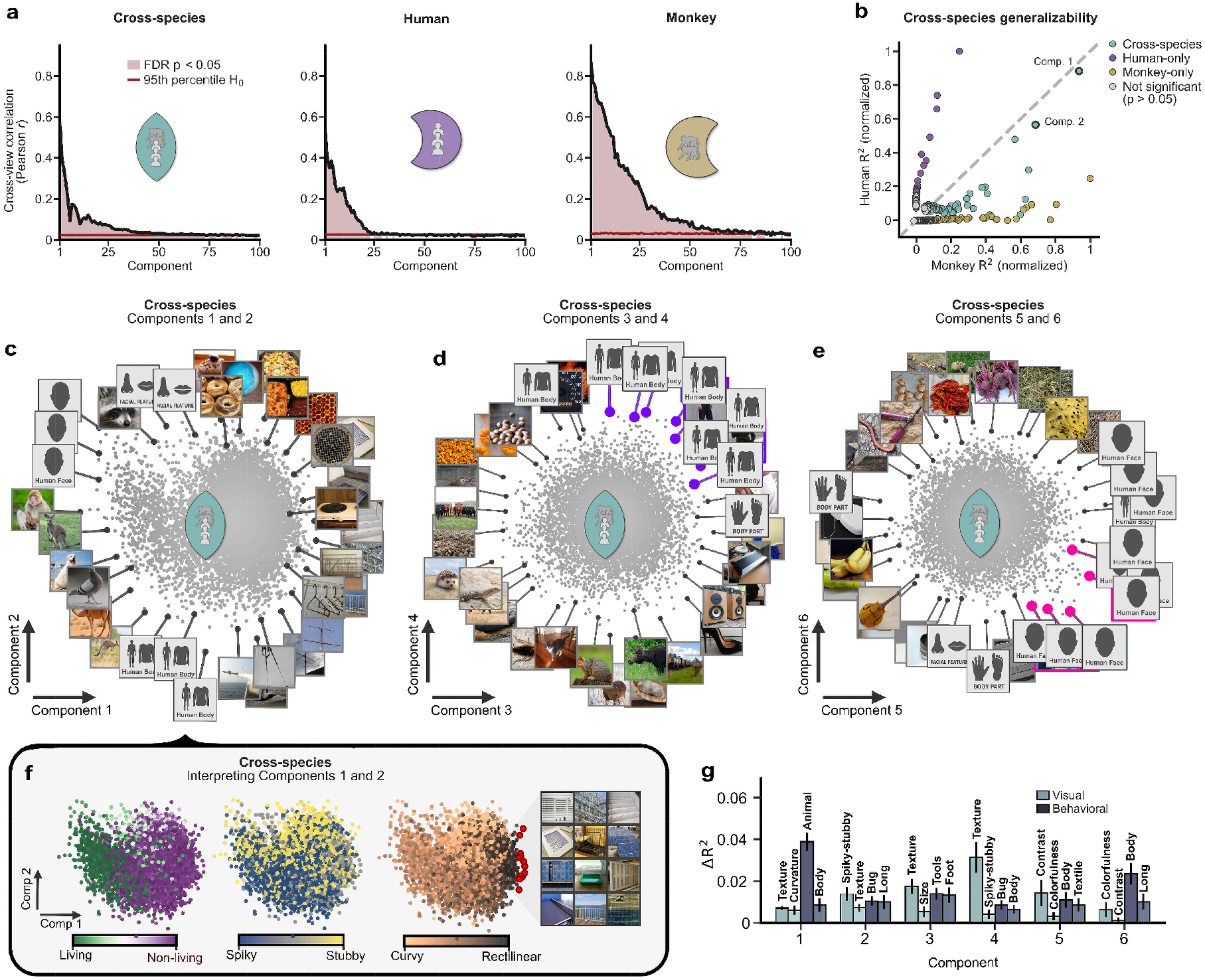
Canonical shared and species-specific spaces of primate IT. (a) Each alignment space shows evidence of high intrinsic dimensionality, indicated by the number of components exceeding the null distribution. (b) Components of the shared space were predicted more evenly from both species, whereas components of the within-species spaces were predicted preferentially from their corresponding species (cross-validated *R*^2^ from native IT responses, normalized within species). The most predictive pair of components of the shared space are labeled “Comp. 1” and “Comp. 2”, and visualized in panels c and f. (c) Image-wise loadings of the first two cross-species axes, with high-loading images displayed to aid interpretability. (d) Same visualization as in c, but for cross-species components C3–C4. Purple callouts highlight a body-silhouette cluster in the upper-right quadrant. (e) Cross-species components C5–C6. Magenta callouts highlight a face-related cluster in the lower-right quadrant. (f) Embeddings for the first two cross-species components as in c, but colored by livingness ratings, spikiness, and rectilinearity. (g) Feature-regression analysis of the first six cross-species components. Bars show the two strongest visual and behavioral predictors for each component (unique contribution: drop in cross-validated *R*^2^ when that predictor is removed from a joint ridge model).

Although the cross-species space was fit jointly to human and macaque data, its components could still be driven disproportionately by structure from one species alone. We therefore asked how well each component could be predicted from native IT response patterns in both species, using cross-validated ridge regression in a leave-one-subject-out scheme (see Methods). We found that shared components were explained more evenly by both species than components derived from the human-only or macaque-only fits, quantified based on each model’s deviation from the diagonal in Fig. 2B (*p <* 0.02 for both models). This demonstrates that relative to the within-species alignments, the alignment between species recovered representational structure more jointly characteristic of both species.

### Similarities across primate species

Having defined the cross-species and within-species spaces, we next asked what structure was captured by the shared alignment. We found that the two leading shared axes coarsely aligned with animacy and spikiness, two dimensions previously proposed to organize object space in primate IT (Kriegeskorte et al., 2008; Bao et al., 2020; Coggan and Tong, 2023). In support of this, the first canonical axis correlated robustly with independently derived livingness ratings (*r* = *−*0.57, *p <* 0.001; Stoinski et al., 2024; Fig. 2F). Exceptions to this pattern were broadly consistent with an animacy interpretation: images of non-living things loading on the animate pole often conveyed animate structure, including puppets, masks, and objects being held or worn by humans (Fig. S6). The second axis captured a continuum from spiky to stubby objects (Fig. 2F). In line with this, the first principal component of AlexNet layer fc6, previously used to operationalize this dimension (Bao et al., 2020), predicted this axis better than all other canonical axes (*R*^2^ = 0.26 *±* 0.03; all pairwise comparisons *t >* 12.75, *p <* 0.001, FDR-corrected; Fig. S7). In support of this finding, a visual feature computing spikiness directly from segmented object masks also emerged as the strongest predictor in the feature-regression analysis (Fig. 2G). These findings show that the alignment recovered established dimensions of IT organization without the use of predefined category labels or feature axes to fit the model. Importantly, the leading axes were not exhaustively explained by animacy and spikiness. Stimuli loading highest on the inanimate side of the first axis contained grid-like rectilinear patterns, in line with the known sensitivity of IT to rectilinear objects (Fig. 2F; Yao et al., 2023; Nasr et al., 2014; Long et al., 2018). The richness of IT organization was also apparent beyond the two leading axes, with subsequent reliable components containing meaningful structure (Fig. 2D–E; Figs. S2–S5). For example, across C3–C4, animals varied in orientation, with horizontally posed animals in the bottom left and vertically oriented bodies in the top right quadrant. Along C5–C6, images with dark or bright empty backgrounds appeared in the lower left quadrant, while richly textured objects and scenes featured in the top right quadrant (Fig. 2D–E). In addition, we observed body- and face-related clusters in these spaces (Fig. 2D, upper right, purple; Fig. 2E, lower right, magenta). These observations were reinforced by analyses showing a broad enrichment of faces and bodies across components (Fig. S8). Thus, beyond the coarse functional organization emphasized in previous work, our findings revealed a much richer representational geometry, with structure demonstrating the presence of further shared representations. These observations motivated a further decomposition of the cross-species space into more interpretable dimensions.

### Parts-based decomposition of the shared space

The canonical spaces provided meaningful axes while also hinting at more local representational structure. To characterize this structure, we used symmetric non-negative matrix factorization (sNMF; Ding et al., 2005; Kuang et al., 2012), which offers a complementary, parts-based decomposition of the stimulus-similarity structure (Khosla et al., 2022; van Dyck et al., 2026). Specifically, we constructed a stimulus-by-stimulus similarity matrix from the cross-species space and factorized it using sNMF (Fig. 3A; see Methods). To identify factors shared across primates, we retained only those whose stimulus weights could be recovered from both the human and macaque spaces, and ranked them by their cross-species generalization. Figure 3B shows the twelve strongest factors, which we interpreted through their highest-weighted stimuli and word clouds quantifying the most strongly enriched basic object categories (see Fig. S11 for a more extensive overview). Under these criteria, individual factors appear to be dominated by interpretable contents (Fig. 3B).

**Fig. 3.**
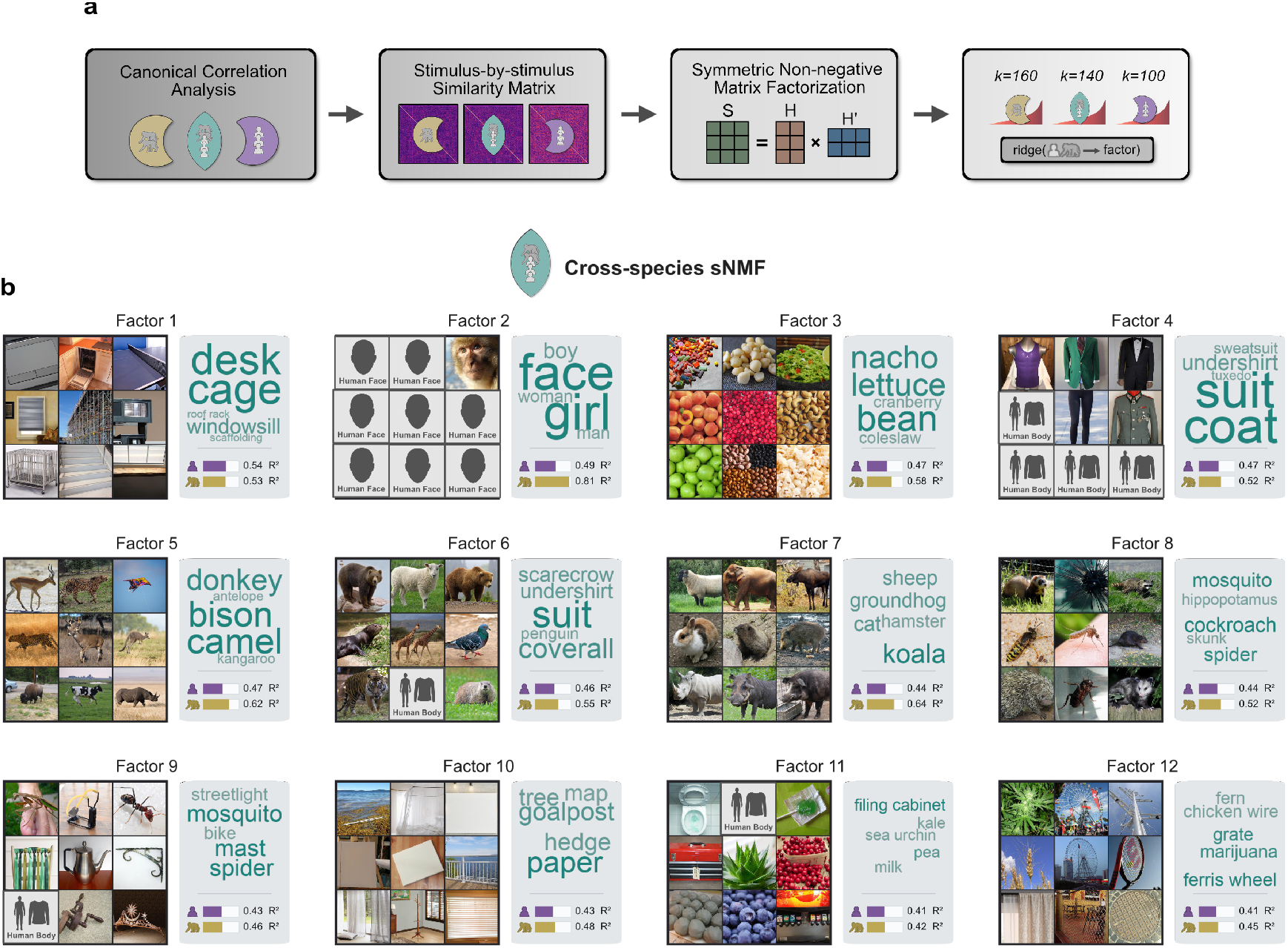
Parts-based factorization of the cross-species space. (a) Overview of the factorization pipeline. We computed stimulus-by-stimulus similarity matrices from the universal and within-species MCCA spaces and factorized them with sNMF. Cross-species factors were retained if recoverable from both species’ projections, whereas within-species factors were retained if recoverable from the corresponding species but substantially less so from the other. Factors were ranked by cross-validated *R*^2^. (b) Twelve most predictive sNMF factors derived from the shared space, shown with top image loadings and word clouds that reflect category enrichments. Inset bars indicate the mean cross-validated ridge *R*^2^ for recovering each factor’s stimulus weights from the human and macaque projections on the cross-species space.

Consistent with analyses over the cross-species space, sNMF recovered factors commonly associated with object categories (Yargholi and Op de Beeck, 2023), such as faces (factor 2) and body silhouettes (factor 4). It also uncovered factors dominated by rectilinear grids (factor 1) and small, granular objects (factor 3), as well as other interpretable dimensions (see Fig. S11 for an overview). Strikingly, several factors captured mixtures of visual and conceptual features—for example, factors 6 and 7 both relayed animate contents, but distinguished right-versus left-facing animals. Similarly, spiky content emerged in both factors 8 and 9, but the former captured animate content, while the latter showed more inanimate objects.

Taken together, these findings demonstrate that the shared organization of primate IT cannot be reduced to a small set of broad axes. Instead, our analyses identify the existence of a rich set of shared dimensions that capture visual properties, salient high-level categories such as biological contents, and specific interactions of these two types of features. The finding that features remain intermixed even across finer decompositions suggests that IT contains conjunctions of visual and concept-level properties (Ratan Murty and Arun, 2018).

### Differences between within-species spaces

Having established commonalities between human and macaque IT in the cross-species space, we next aimed to identify differences between the human and macaque within-species spaces. We examined these differences with the premise that a direct contrast between spaces can reveal dimensions or features differentially expressed in human fMRI patterns and macaque recordings. Across approaches from different angles, we found converging evidence for an imbalance in the representation of animacy, a conspecific preference, and differences in the relative contribution of visual and conceptual information to these spaces.

### Differences in basic object categories

First, we asked which of the 720 basic object categories in our stimulus set (Fig. 1) were more strongly expressed in either the human or macaque space. We quantified each category’s relative expression in the human and macaque spaces, using the cross-species space as a baseline. Their difference defined a species-asymmetry index (*z*_Human_ *− z*_Monkey_), with positive values indicating stronger expression in the human space and negative values stronger expression in the macaque space. This analysis revealed a striking animacy asymmetry: the macaque space was biased toward living categories, whereas the human space was biased toward non-living categories (Fig. 4A–B; *r* = *−*0.381, *p* = 0.001). Despite the differences in measurement modality between humans and macaques, this result suggests that animate stimuli carry greater representational salience in macaque IT, whereas human artifacts are more strongly represented in human IT. More broadly, this indicates that species-specific ecological and behavioral priorities are embedded already within the representational organization of IT.

**Fig. 4.**
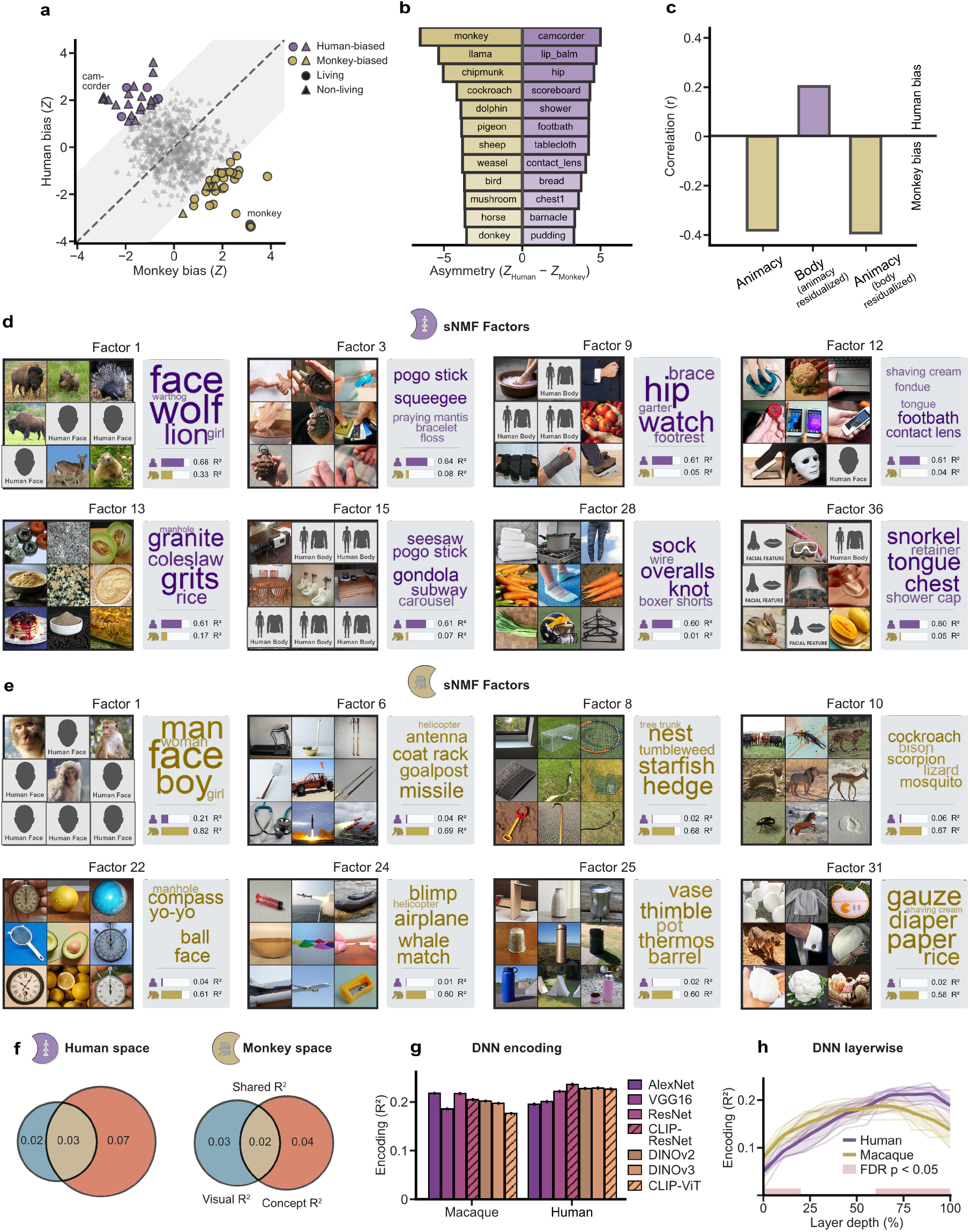
Differences between human and macaque canonical spaces. (a) Species-asymmetry index for each of the 720 basic object categories. Positive values indicate stronger expression in the human space and negative values stronger expression in the macaque space. Points outside the permutation-derived null band indicate categories with significant asymmetry scores, and marker shape denotes living versus non-living categories. (b) Basic categories with the strongest macaque-relative expression (left) and human-relative expression (right), ranked by species asymmetry. (c) Correlation of species asymmetry with animacy, human body-part content after controlling for animacy, and animacy after controlling for body-part content. (d) Example human-specific sNMF factors from the within-species spaces, shown with top-loading images and category-enrichment word clouds. (e) Macaque-specific sNMF factors. Within-species factors were retained if they were recoverable from the matching species CCA space and substantially weaker in the other species (indicated by inset bars). (f) Visual and concept-level variance partitioning in each species space: Venn diagram shows unique and shared explained variance. (g) Best-layer encoding performance across DNNs for macaque and human spaces. (h) Layer-wise DNN encoding across models; thin lines show individual models, thick lines show species means, and red segments mark layer depths with statistically robust species differences.

We further observed that many human-biased categories were rich with human body parts. For instance, nearly half of the camcorder images included hands, and all but one of the footbath examples contained feet. This raised the possibility that the animacy asymmetry was alternatively driven by human effectors appearing alongside, or interacting with, otherwise inanimate objects, such as manmade tools. However, the animacy asymmetry persisted after controlling for a behavior-derived body-part dimension (Hebart et al., 2023) (Fig. 4C; partial *r* = *−*0.392, *p* = 0.001, dimension displayed in Fig. S9), while body-part content retained a smaller independent association with species asymmetry (partial *r* = 0.201, *p* = 0.001). Together, these results point to two separable sources of asymmetry between each primate space: a broad imbalance in the relative salience of animate and inanimate content across the two spaces, and a human conspecific bias for human body parts.

### Species-specific factorization

Moving beyond coarse category-level imbalances, we next asked which representational dimensions differentiated the two within-species spaces. We factorized each space separately, retaining factors that generalized within species but only weakly to the other species’ space (see Methods). Several human-space factors captured human body parts, as evidenced by their top images and category enrichments (Fig. 4D; factors 3, 9 and 12). Consistent with this interpretation, the behavior-derived body-parts dimension introduced earlier predicted two human-specific factors more strongly than any macaque-specific factor (*R*^2^ = 0.034–0.042; both FDR-corrected *p <* 0.001; Fig. S10). Conversely, the most predictive macaque-space factor included monkeys among its top-loading images, consistent with “monkey” being the most strongly expressed category in the macaque space (Fig. 4B). These results support the interpretation that a key difference across these primate spaces is the salience associated with conspecific stimulus materials—such as human hands and body parts for humans, and images containing monkeys for macaques. Separately, these within-species factors further showed that visual and conceptual features were not cleanly separable: factor 31 emphasized stimuli that were white in color and stubby in shape, while factors 24 and 25 emphasized horizontally and vertically oriented inanimate spiky objects, respectively (see Fig. S12 and S13 for a more extensive overview of within-species factors).

### Differences in visual versus conceptual features

Given that human visual representations are thought to be shaped not only by visual experience but also by object-specific semantic associations expected to be weaker or absent in macaque monkeys, we asked whether the relative contributions of visual and conceptual information differed between the two species-specific spaces. To this end, we predicted the MCCA components of each space with ridge regression models that included only visual features, only concept-level features, and a joint model including both (Fig. 4F). Visual predictors spanned low-level (contrast, texture, spatial frequency) and mid-level properties (spikiness, real-world size; Konkle and Oliva, 2012), whereas conceptual predictors indexed category membership (e.g., mammal, electronic device, plant). In the human space, concept-level predictors explained substantially more unique variance than visual predictors (concept *R*^2^ = 0.071 vs visual *R*^2^ = 0.022; Fig. 4F; *p <* 0.001). In contrast, the macaque space was more balanced (concept *R*^2^ = 0.042 vs visual *R*^2^ = 0.032; *p <* 0.001). These findings reveal a conceptual-perceptual asymmetry between the canonical spaces, with the human space more strongly predicted by high-level conceptual over visual information relative to the macaque space (Li et al., 2026).

Finally, we asked whether this asymmetry was also reflected across the processing hierarchies of deep neural networks (DNNs). We reasoned that spaces emphasizing relatively lower-level visual information should be predicted better by earlier network layers, whereas spaces emphasizing more integrated object-level information should favor later layers. To evaluate this, we predicted each space using layer-wise activations from seven networks. Although these DNNs varied in objective function, architecture, and training set, all seven networks achieved similar predictive performance for the canonical spaces at their optimal layers (*R*^2^ = 0.196–0.236 for humans and 0.177–0.218 for macaques; Fig. 4G; Conwell et al., 2025). In support of an asymmetry, the earliest depth bin (0–20%) predicted the macaque canonical space better (FDR-adjusted *p* = 0.012), whereas the two deepest bins (60–80% and 80–100%) predicted the human space better (FDR-adjusted *p* = 0.002 and *p <* 0.001; Fig. 4H). This layer-wise dissociation provides complementary evidence that the human space is relatively more strongly associated with higher-level information.

### Orthography-related structure in shared IT space

Whilst exploring the alignment between species, we noticed that the top-weighted images for some shared components, such as component 7, contained symbols (e.g., letters and numbers), icons (e.g., hearts and crosses), and ensembles of multiple objects (e.g., arrays of tools; Fig. 5A). This shared structure is surprising, given that many of these stimuli are human signifiers with no clear meaning and no obvious significance to macaque monkeys. This raises the possibility that the cross-species space captured a shared neural signature that in macaques may constitute a precursor to orthographic representations (Dehaene and Cohen, 2007). To test this hypothesis without decoding directly from the space we had inspected, we refit the CCA within each cross-validation fold and decoded labels for symbols, icons, and multi-object ensembles from held-out images, separately for the human and macaque projections onto the shared space. As positive and negative controls, we included reference decoders based on the strongly shared representations of faces and bodies, as well as a random image set baseline, respectively.

**Fig. 5.**
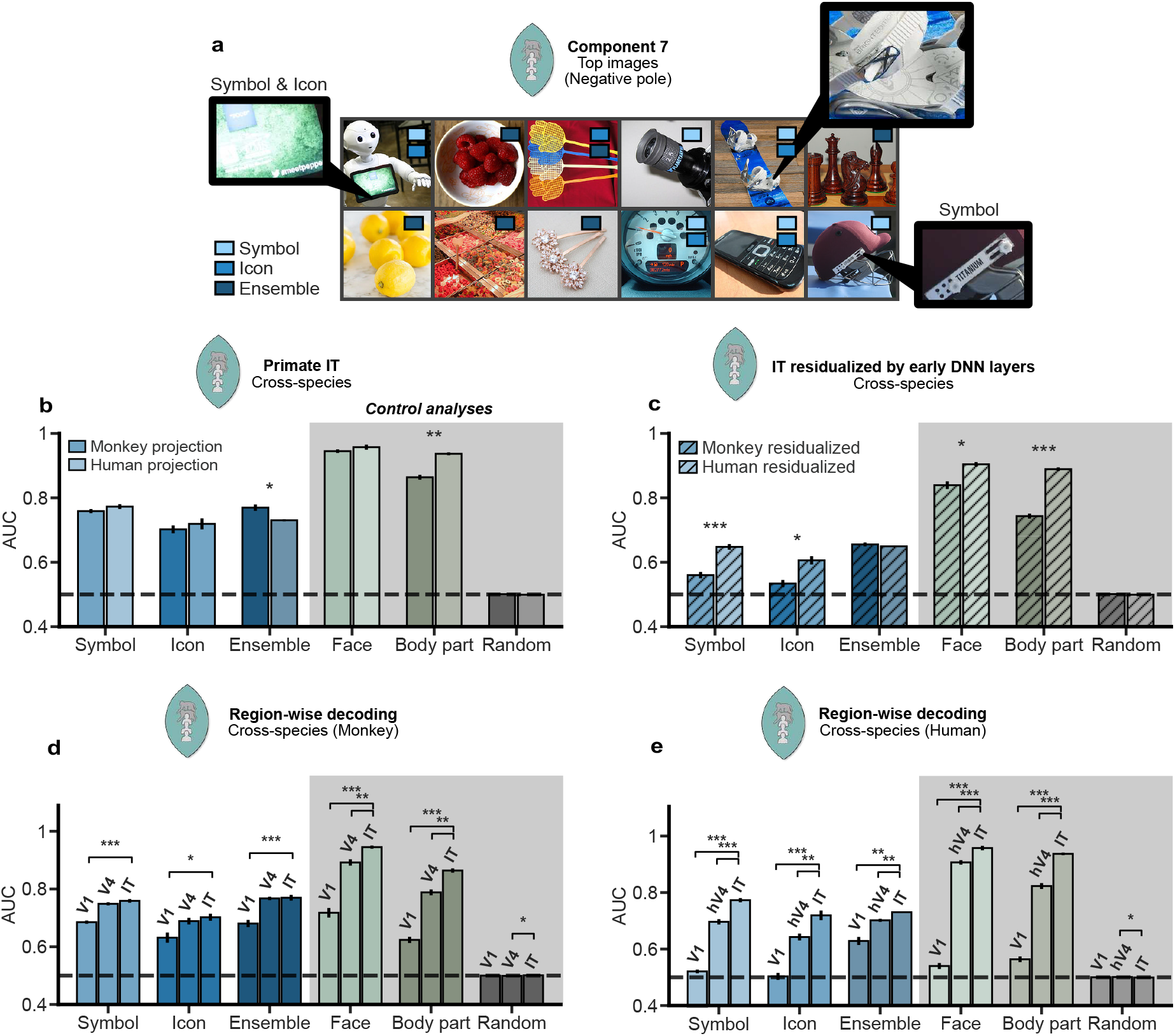
Orthography-related structure in the shared primate IT space. (a) Top-loading images from the negative pole of universal CCA component 7; colored markers inside the images denote the presence of symbols, icons, and multi-object ensembles. (b) Cross-validated logistic-regression decoding (area under curve; AUC) of symbol, icon, and ensemble labels from macaque versus human projections (i.e., views) onto the shared MCCA space; face, body-part, and random image set decoders serve as reference analyses. (c) Decoding after residualizing each species’ projections onto the universal space by regressing out a control embedding derived from aligned early-layer activations across seven DNNs. (d) Region-wise decoding across macaque projections onto universal CCA spaces fit separately in V1, V4, and IT. (e) Same as (d), but using human projections (and area hV4 instead of V4). Error bars indicate SEM across folds; dashed line indicates chance (AUC = 0.5); significance markers denote FDR-adjusted comparisons *p <* 0.05 (*), *p <* 0.01 (**), and *p <* 0.001 (***).

We found that the shared IT space contained robustly decodable information for symbols, icons, and multi-object ensembles, with reliable performance in both macaques and humans (all FDR-adjusted *p* = 0.005; permutation test; Fig. 5B). To evaluate whether this was explained by low-level properties alone, we residualized the primate projections by regressing out a control embedding obtained by aligning early-layer activations across the seven DNNs introduced above. Factoring out this variance strongly reduced decoding performance for all orthography-related decoders in both species (reductions of 0.08–0.20 AUC; all FDR-adjusted *p <* 0.001), yet decoding remained reliably above chance for every label in both species (FDR-adjusted *p ≤* 0.015; Fig. 5C). After residualization, human decoding exceeded macaque decoding for symbols and icons (ΔAUC_human_*_−_*_macaque_ = 0.088 and 0.072; FDR-adjusted *p <* 0.011), whereas ensemble decoding did not differ reliably between species (ΔAUC_human_*_−_*_macaque_ = *−*0.006; FDR-adjusted *p* = 0.309).

Lastly, to establish whether evidence for this shared representational structure was already present in earlier visual areas, we repeated the analysis using recordings from V1 and V4/hV4 across these datasets. Orthographic decoding increased from V1 to IT in both species (all IT–V1 contrasts, *p <* 0.02; Fig. 5D–E). From V4/hV4 to IT, humans showed reliable gains for symbols, icons, and ensembles (ΔAUC = 0.029–0.077; all FDR-adjusted *p <* 0.005), whereas macaques did not (ΔAUC = 0.002–0.013; all FDR-adjusted *p ≥* 0.055). Directly comparing these gains across species, the V4-to-IT increase was larger in humans than in macaques for symbols and icons (ΔAUC = 0.066 and 0.063; FDR-adjusted *p ≤* 0.007), but not for ensembles (*p* = 0.052). Together, these analyses provide converging evidence that stimulus contents broadly related to orthography are structurally aligned across macaque and human visual cortex, and that low-level visual properties account for part, but not all, of this alignment. In addition, humans showed stronger residual decoding for symbols and icons and a steeper V4-to-IT gain across orthographic features, possibly reflecting additional specialization or abstraction in human ventral visual cortex.

## Discussion

Here, we systematically examined the similarities and differences in the functional organization of primate inferotemporal cortex (IT) using human fMRI and macaque intracranial responses to a large and diverse set of naturalistic object images. By aligning these responses using a multivariate data-driven framework, we uncovered a rich, high-dimensional representational space shared between species, demonstrating that the shared representational geometry extends well beyond a few established dimensions. A comparison of species-specific spaces further allowed us to identify divergent measured representations, revealing a stronger preference for high-level, conceptual content in humans as compared to macaques, and with both species carrying a conspecificity preference. The approach also uncovered surprising similarities, including a shared neural sensitivity to orthographic materials, suggesting that representations often thought to be uniquely human may build on precursors present in macaque IT. Together, these findings clarify the extent to which IT organization generalizes across primates, revealing a rich shared geometry while identifying dimensions along which humans and macaques diverge.

Previous work comparing human and macaque IT has established functional correspondences along several prominent dimensions, including animacy, shape, and other mid-level features (Kriegeskorte et al., 2008; Bao et al., 2020; Yao et al., 2023). Recent work has further shown representational alignment across scene stimuli (Vinken et al., 2025; Li et al., 2026). However, it remains unclear how cross-species functional organization generalizes across object space more broadly, including whether similarities are confined to a small set of dominant axes or whether there is a broader geometry shared across species. Here, we show that human and macaque IT indeed share a rich, multidimensional organization of which established dimensions form only a limited part. This finding extends high-dimensional accounts of IT organization within individual species (Lehky et al., 2014; Gauthaman et al., 2025; Zhang et al., 2026) by showing that much of this representational complexity is shared across species, placing cross-species inference on firmer ground. A closer examination of the aligned space revealed leading dimensions broadly corresponding to animacy and spikiness, even though neither dimension was specified in advance. However, the first axis also carried structure related to rectilinearity, showing that these axis labels provide only partial descriptions of the underlying organization. Later axes revealed diverse interpretable structure across the shared geometry, including animate contents varying in orientation and local face and body clusters spanning multiple axes. Factorizing the space as a whole yielded dimensions spanning low-level visual properties, shape and orientation, biologically salient content, and concept-level organization. Several factors captured specific conjunctions of visual and conceptual properties—for example, animacy together with orientation or spikiness together with object category—suggesting that IT jointly organizes visual and conceptual information.

Beyond commonalities, the within-species contrasts revealed differences at the level of object categories, in the factorized dimensions we identified, and in the encoding of visual versus concept-level information. Across these analyses, a recurring pattern was that each species appeared to emphasize conspecific contents: human body parts were enriched in the human space, and stimuli containing macaques and other monkey taxa loaded more strongly onto the macaque space. For humans, this was evident in the stronger expression of categories featuring human effectors (Fig. 4C), the dominance of human body parts in human-selective sNMF factors (Fig. 4D), and their broad enrichment among the top loadings of the human space (Fig. S8). Conversely, the monkey stimulus category, composed of macaques and other monkey taxa, showed the strongest macaque-over-human bias across all 720 categories (Fig. 4B).

The datasets differed in recording modality (human fMRI versus macaque multi-unit activity), anatomical sampling, and task parameters, meaning species and methodological effects cannot be fully disentangled. In spite of this, the cross-species alignment recovered many reliably shared axes, including established dimensions of animacy and spikiness, suggesting that the analytical approach is able to discover shared representations of primate IT even in the face of measurement differences. Within-species contrasts warrant greater caution because effects between the spaces may partially reflect dataset-specific structure. Nevertheless, we find several theoretically plausible results that are unlikely to be explained primarily by methodological variation: Across analyses, we find an emphasis on human body parts in the human space, monkeys in the macaque space, and stronger conceptual prediction of human IT. Finally, one potential limitation applicable more broadly to the field is that our conceptual predictors and DNN embeddings might contain human biases; later DNN layers, for example, may favor the human space because the networks were trained on human-curated datasets. Future work could address these biases by using stimulus sets more reflective of macaque visual experience, or by estimating relevant distinctions in the visual world through macaque behavior (Cherian et al., 2026).

The spaces computed by this approach can also be used to form hypotheses which can be tested quantitatively. When inspecting the cross-species space, we found several components with images containing symbols, icons, and multi-object ensembles, raising the possibility that orthography-related content has a shared representational structure in primate IT. Previous work has shown specialized responses to orthographic stimuli in human ventral cortex (Cohen et al., 2000), sensitivity to symbols in macaque IT (Rajalingham et al., 2020; Srihasam et al., 2012), and numerosity sensitivity in macaque parietal and prefrontal cortex under ensembles of geometric elements (Viswanathan and Nieder, 2013). Here, we show that related structure is not confined to either species alone, but can be decoded from a linear subspace spanning human and macaque IT. These results are consistent with the neural recycling hypothesis, which postulates that human orthographic capacities build on ventral-stream mechanisms already present in other primates (Dehaene and Cohen, 2007).

How do these results inform theoretical accounts on the nature of IT representations? One interpretation is that primate IT is formatted by continuous axes (Bao et al., 2020; Coggan and Tong, 2023; Yao et al., 2023; Lehky et al., 2014; Gauthaman et al., 2025). An alternative interpretation is that primate IT is organized by categories—a distinct format that registers discrete classes of content such as faces, bodies and other salient contents as the primary unit (Kiani et al., 2007; Yargholi and Op de Beeck, 2023). A more recent study suggests a third possibility, namely that the representational format itself changes dynamically (Shi et al., 2026). Our results show a mixture of evidence: we find a multidimensional space of axes, with category-related clustering within that space. Faces are enriched across cross-species axes, and sNMF separates faces, body parts, body silhouettes, and animals into distinct groupings. These observations suggest two possible reconciliations. On the one hand, categories and continuous axes may coexist as two representational formats in primate IT, with different analytical tools emphasizing one or the other. For example, methods such as CCA or principal component analysis may probe graded representations, and non-negative, parts-based methods such as sNMF may emphasize categories. On the other hand, IT may have a single underlying format, with different decomposition methods yielding summaries that make the same representations appear more category- or dimension-like (van Dyck et al., 2026). Adjudicating between these possibilities requires identifying under which conditions the two formats make different empirical predictions, for example using computational models.

The alignment framework put forward in the present work extends beyond cross-species comparisons, offering a general strategy for mapping shared and distinct representations across any domains with repeated measurements. In this study, these domains were species, the measurements were individual subjects, and the explanatory target was functional architecture. But the logic extends to comparisons between neuroimaging modalities, behavioral tasks, brain regions, and temporal windows. In each case, cross-domain alignments isolate structure that generalizes, while contrasts between within-domain alignments show what is differentially expressed. Thus, this framework is not an algorithmic innovation but a way to harness existing alignment algorithms for studying cross-domain similarities and differences. Here, we used MCCA because of desirable properties such as its view-wise projections (Fig. 5), but the same approach could be implemented with shared response modeling (Chen et al., 2015) or other alignment techniques (Haxby et al., 2011; Chen et al., 2014).

## Acknowledgments

We thank Paolo Papale for acquiring and preparing the monkey dataset and Florian Mahner for his advice and methods code. We thank Chris Baker, Kurt Braunlich, Jonas Noëlle, Luca Kolibius, Sophia Snipes and the Hebart lab for useful discussions on this work.

## Author contributions

Conceptualization: SVB, MNH

Methodology: SVB

Investigation: SVB

Visualization: SVB

Funding acquisition: MNH

Supervision: MNH

Writing – original draft: SVB

Writing – review & editing: SVB, MNH

## Data and materials availability

The human THINGS-fMRI single-trial response data are publicly available through Figshare+ (doi:10.25452/figshare.plus.c.6161151), and the macaque THINGS Ventral-stream Spiking Dataset is publicly available through G-Node GIN (doi:10.12751/g-node.hc7zlv). Newly derived image properties and all analysis code will be made available to editors and reviewers during peer review and deposited on GitHub by the time of publication.

## Methods

### Stimuli and datasets

#### Stimuli

We used naturalistic object images from the THINGS database (Hebart et al., 2019). We restricted all analyses to the intersection of stimuli shared across the macaque and human dataset, yielding 8,640 images from 720 basic object categories (12 exemplars each).

#### Macaque electrophysiology

We used multi-unit activity from the THINGS Ventral Stream Spiking Dataset (TVSD; Papale et al., 2025), which comprises 1,024-channel recordings from V1, V4, and inferotemporal cortex (IT) in two macaques (N and F). We used the normalized, time-averaged response matrices provided with the dataset and limited all analyses to the shared stimulus set described above. In the TVSD preprocessing pipeline, responses were averaged within ROI-specific post-stimulus windows (V1: 25-125 ms, V4: 50-150 ms, IT: 75-175 ms). Unless otherwise noted, analyses focused on IT responses (75-175 ms). We confirmed using an independent temporal generalization analysis that this window captures stable category-level structure (Fig. S1). We retained channels with a non-negative reliability (cutoff *≥* 0), defined as the average pairwise correlation of that channel’s responses across 30 repeated presentations of 100 test images, resulting in 250 IT channels for monkey N and 313 for monkey F. Full experimental details, including implant locations, are provided in Papale et al. (2025).

#### Human fMRI

We used human fMRI data from THINGS-data (Hebart et al., 2023; Contier et al., 2024), collected from three participants across multiple sessions. In this experiment, images were presented for 500 ms followed by 4 s of fixation. Image-wise response estimates were obtained using single-trial models with voxel-wise hemodynamic response function estimation (Contier et al., 2024). Full acquisition and preprocessing details are described in Contier et al. (2024) and Hebart et al. (2023). For the main analyses, we defined an IT region of interest (ROI) as the union of voxels in lateral occipital complex (LOC), fusiform face area (FFA), parahippocampal place area (PPA), extrastriate body area (EBA), and pRF-defined LO1 and LO2. Similarly to the macaque channels, we retained voxels with a reliability of *≥* 0, yielding 3,929, 3,187, and 2,309 IT voxels for human participants 1-3 (Fig. S1). For region-wise control analyses in the orthographic analyses, V1 and hV4 were defined using the same voxel metadata.

#### View construction and preprocessing

We treated each of the five subjects (two macaques and three humans) as an independent view for crossspecies alignment. For each view, we constructed a stimulus-by-feature matrix (N stimuli x P features), where features corresponded to macaque recording channels or human fMRI voxels. To reduce session- and run-specific baseline differences in the human data, voxel responses were mean-centered within each session-run block while preserving stimulus-driven variance. All features were z-scored per view within each cross-validation fold, using means and standard deviations estimated from the training set. Then, we applied a series of decomposition techniques, such as multiset canonical correlation analysis (MCCA) and non-negative matrix factorization (NMF) over MCCA space. Throughout this manuscript, we refer to CCA-derived embeddings as components (or axes), NMF-derived embeddings as factors, and dimensions as an umbrella term.

### Multiset canonical correlation analysis

We aligned neural responses across subjects using regularized MCCA (Hardoon et al., 2004; Correa et al., 2010; de Cheveigné et al., 2019). For each view *X*^(^*^v^*^)^, MCCA estimates a linear projection *W* ^(^*^v^*^)^, yielding canonical scores *Y* ^(^*^v^*^)^ = *X*^(^*^v^*^)^*W* ^(^*^v^*^)^ that maximize shared correlation across views. Specifically, regularized MCCA finds a shared canonical coordinate space by solving a generalized eigenvalue problem on block covariance matrices composed of within-view and between-view covariances. To ensure a stable solution, a ridge penalty is applied to shrink the projection weights (for details, see Bilenko and Gallant, 2016).

We fit three model families. The universal model was fit across all five views (two macaques and three humans) and was used to isolate structure shared across primates. In addition, we fit a human-only model across the three human views and a macaque-only model across the two macaque views to characterize structure reproducible within each species.

Regularization strength (*λ*) was selected separately for each family by nested cross-validation on held-out stimuli. Specifically, we evaluated 15 log-spaced ridge values (*λ ∈* [10^1^, 10^7^]) with repeated nested cross-validation (three repetitions; outer 5-fold, inner 3-fold). Within each split, we selected the *λ* value that maximized mean held-out cross-view correlation. This procedure yielded *λ* = 10^4^ for the universal and human-only families and *λ* = 10^2^.^7^ for the macaque-only family (Fig. S14).

#### Canonical spaces

Using the selected regularization for each family, we fit a final MCCA model (k = 100) and evaluated the held-out cross-view correlation of each component. To establish which components are statistically reliable and suited for subsequent analyses, we generated a null distribution by repeating the same MCCA fit with permuted stimulus labels (200 permutations). This abolishes stimulus-dependent latent structure common across subjects, yielding expected cross-view correlations under the null hypothesis.

For each component, we computed a *p*-value as the fraction of null correlations exceeding the empirical correlation derived from true labels, adjusting the *p*-values using Benjamini-Hochberg false discovery rate (FDR) correction (Benjamini and Hochberg, 1995). For all downstream analyses, we used only the subset of components with an FDR-adjusted *p <* 0.05. These retained MCCA components reflect linear transformations that reliably align subjects, and throughout this manuscript we refer to retained components as a family’s “canonical space”. All CCA analyses were implemented using Pyrcca (Bilenko and Gallant, 2016).

When a single embedding was required for downstream analyses, we averaged view-specific canonical scores within a model family to obtain one score per stimulus and component. In the orthographic decoding analyses, where we compared how the universal space was expressed in humans and macaques, we standardized universal-space component scores within each view and then averaged canonical scores within species rather than across all five views.

#### Recoverability from IT

To quantify how strongly each MCCA component was supported by humans versus macaques, we used ridge regression to predict family-level component scores from subjects’ original channels or voxels, averaging prediction performance within species. This offers a flexible way to quantify to what extent any axis is recoverable from native IT, and driven by one versus another species. For a given subject, targets were defined in a leave-one-view-out manner by averaging component scores across the remaining views in the same model family. Because each target excluded the held-out subject’s own view, this analysis indexed support for family-level axes rather than reconstruction of the subject’s own canonical scores. Predictors were standardized within each training fold. To prevent predictions from being driven by the varying dimensionality of each subject’s data, we first reduced each native space to 250 PCA components, matching the subject with the fewest features (monkey N). Ridge performance was summarized as cross-validated *R*^2^ per component, and in Fig. 2B, normalized within each species by dividing by the maximum *R*^2^ across all model families, placing both species on a common scale for visualization. To summarize species balance, we computed a per-component imbalance score as the squared difference between normalized human and macaque recoverability 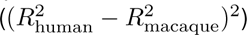, and averaged this value across FDR-retained components within each model family. We compared mean imbalance between the cross-species family and each within-species family using one-sided component-level permutation tests that shuffled family labels while preserving the number of retained components per family.

### Feature-based characterization of canonical spaces

#### Feature banks

To characterize what information the canonical spaces encode, we assembled three complementary sets of stimulus descriptors. The first set comprised seven visual properties spanning low- and mid-level image statistics. Five of these were computed directly from images: RMS contrast, colorfulness (a composite of red-green and yellow-blue channel variances), spatial frequency (variance of the Laplacian), texture (mean Sobel gradient magnitude), and spikiness (the inverse ratio of object area to convex-hull area, computed from segmented object masks). Curvature and real-world size were obtained from human ratings, per-image for curvature, and for every THINGS category for real-world size (THINGSplus; Stoinski et al., 2024). All seven properties were rank-normalized to the unit interval after applying a log transform to features with skewness *>* 1 or kurtosis *>* 3.

The second set comprised 66 behavioral dimensions from the SPoSE embedding, a sparse non-negative embedding of human object similarity judgments derived from *∼*4.7 million odd-one-out triplet responses (Hebart et al., 2023). Each dimension is interpretable and captures a mixture of perceptual and conceptual properties (e.g., animal-related, red, metallic/artificial). Because the embedding is defined at the concept level, all 12 exemplars of a category received the same 66-dimensional score vector.

The third set comprised 53 binary conceptual category attributes from the THINGS taxonomy (e.g., animal, bird, furniture, vehicle; Stoinski et al., 2024; Hebart et al., 2019), indicating whether each concept belonged to a given superordinate category. These three feature sets were used in complementary ways: visual and behavioral features jointly for characterizing individual canonical components (below), and visual versus concept-level features for contrasting species spaces (see “Comparisons between human and macaque canonical spaces”).

#### Component-wise feature regression

To identify which features drive each canonical axis, we predicted component scores from the joint set of 7 visual properties and 66 behavioral dimensions using ridge regression. Regularization strength was selected per component from 11 log-spaced values (*α ∈* [10*^−^*^2^, 10^6^]) via 3-fold inner cross-validation. Prediction performance was evaluated using repeated *k*-fold cross-validation (5 folds *×* 3 repeats; all features standardized within each training fold). Predictor importance was quantified as the drop in *R*^2^ when removing each predictor from the full model, averaged across outer folds. This analysis was applied to all three model families (universal, human-only, and macaque-only). The main feature-regression panel for the dominant cross-species components is shown in Fig. 2G, with corresponding summaries across model families in Fig. S2C. When summarizing results across components, each component’s contribution was weighted by its held-out cross-view correlation.

#### Top-loading images and cumulative capture curves

We examined component content by ranking stimuli according to their family-averaged canonical scores and inspecting the top-loading images at the negative and positive poles. To summarize whether particular stimulus classes were disproportionately represented among strongly expressed canonical dimensions, we ranked stimuli within each of the first 10 components by the absolute value of their family-averaged canonical scores. For labels of interest, such as faces or body parts, we then computed cumulative capture curves showing the fraction of labeled images recovered as progressively larger fractions of the top-loading stimuli were included (Fig. S8). To summarize how strongly each stimulus was expressed across a canonical space as a whole, we z-scored each of the first 10 components across stimuli, took the absolute value, and averaged across components, yielding one overall loading value per stimulus (Fig. S8).

#### Animacy and spikiness

To relate components to animacy, we correlated image-wise component scores with independently obtained livingness ratings from the THINGS property norms (Stoinski et al., 2024). To assess correspondence with the spikiness axis described by Bao et al. (2020), we extracted AlexNet fc6 activations for the 8,640 stimuli used in this study, rank-transformed each feature across stimuli, computed the first principal component, and used this score to predict component loadings with cross-validated ridge regression. See DNN layer-wise encoding for details on DNN feature extraction.

### Comparisons between human and macaque canonical spaces

#### Category-level species asymmetry

To quantify category-level differences between the human-only and macaque-only spaces, we defined each stimulus’s expression as the L1 norm of its component score vector and averaged this value across the 12 exemplars of each category. Category expression in each species was normalized by the corresponding value in the universal space, z-scored within species, and converted to an asymmetry score defined as human minus macaque. Null bands were obtained by permuting stimulus identities 1,000 times. We related asymmetry scores to THINGS livingness ratings and to a SPoSE-derived body-part dimension using zero-order and partial correlations. Significance was assessed by permutation tests (1,000 shuffles), and uncertainty was summarized with bootstrap confidence intervals (1,000 resamples).

#### Visual-versus-concept model comparisons

To ask whether the species-specific spaces were better explained by visual or conceptual information, we separately fit ridge models with visual predictors only, concept predictors only, and both together. Unique visual variance was computed as 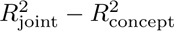, unique concept-level variance as 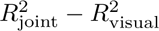, and shared variance as 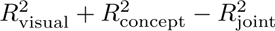 Variance partitioning was summarized across all retained components of each canonical space (32 human and 91 macaque components; Fig. 4F), with component contributions weighted by their held-out cross-view reliability. For statistical inference, both spaces were restricted to their first 32 retained components to match dimensionality and prevent differences in the number of retained components from driving the comparison. Confidence intervals and (p)-values were obtained using 1,000 stimulus-bootstrap resamples.

#### DNN layer-wise encoding

Canonical scores were also predicted from layer-wise activations of seven deep neural networks spanning supervised, contrastive vision-language, and self-supervised objectives: AlexNet, VGG-16, ResNet-50, CLIP-ViT-B/16, CLIP-RN50, DINOv2-ViT-B/14, and DINOv3-ViT-B/16 (Krizhevsky et al., 2017; Simonyan and Zisserman, 2014; He et al., 2016; Radford et al., 2021; Oquab et al., 2024; Siméoni et al., 2026). Input images were resized to a shortest edge of 256 pixels and center-cropped to 224 x 224 pixels. Convolutional feature maps were reduced to one vector per image by global average pooling across spatial dimensions. Transformer activations were pooled according to model family: DINO intermediate-block activations were averaged across patch tokens, with the CLS token used at the final normalized output, whereas CLIP-ViT activations were represented by the CLS token at each sampled block. These layer-wise image vectors were entered into cross-validated ridge models; as in the native-space analysis, the dimensionality of DNN activations was capped at 250 PCA components. Encoding performance was quantified as held-out *R*^2^ for each retained canonical component and summarized as a weighted mean, with component weights proportional to their held-out cross-view correlations. To compare across architectures, layer depth was normalized to the unit interval within each model, and species differences were assessed within normalized depth bins.

### Parts-based decomposition with symmetric NMF

To complement the signed canonical axes with a parts-based perspective, we decomposed stimulus-by-stimulus similarity structure using symmetric non-negative matrix factorization (sNMF; Ding et al., 2005; Kuang et al., 2012). For each model family, cosine-similarity matrices were computed separately per view in canonical-score space, Fisher-z averaged across views, rescaled to the [0,1] interval, and treated as the input similarity matrix for factorization. Factorization rank was selected by matrix-completion cross-validation using an sNMF approach that completes held-out entries (Mahner et al., 2026). Specifically, we repeated the cross-validation procedure five times. In each split, 70% of matrix entries were held out and the factorization was fit on the remaining 30%; the rank minimizing reconstruction error on the held-out 70% was selected. We searched over a rank grid of *r ∈ {*20, 40*,…,* 200*}*. The selected rank was then used to fit the final factorization on the full similarity matrix (*k* = 160, 140, 100, for the macaque, cross-species, and human space, respectively).

To distinguish shared from species-specific factors, each factor’s stimulus weights were predicted from canonical scores via ridge regression with 5-fold cross-validation. In the universal decomposition, factors were labeled shared if they were predictable from both species’ projections (*R*^2^ *≥* 0.20 for each). In within-species decompositions, factors were labeled species-specific if they were predictable from the matching species (*R*^2^ *≥* 0.20) and predictability from the other species was less than half as large 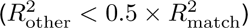 Shared factors were ranked by the minimum of the two species’ cross-validated *R*^2^ values, and species-specific factors were ranked by the matching species’ cross-validated *R*^2^. Factor content was summarized with AUROC-based category enrichment.

To quantify body-part encoding, we predicted each retained within-species factor from the SPoSE body-part dimension using 10-fold cross-validated ridge regression. Significance was assessed by permuting body-part scores across concepts while keeping the 12 exemplars of each basic object together (10,000 permutations), followed by FDR correction across all retained human and macaque factors.

### Orthography-related decoding

To test whether the universal IT space captured aligned structure related to orthographic content, we decoded binary image annotations indicating the presence of symbols, icons, and multi-object ensembles. Face and body-part labels served as positive controls, and random binary labels with approximately 50% prevalence served as negative controls. Images were annotated by visual inspection (S.B.). Symbols were defined as non-pictorial signs, such as letters and numbers; icons as signs referring to real-world objects, such as hearts and arrows; and ensemble images as arrangements containing multiple similar objects. Face and body-part labels included both human and non-human animals. This yielded 1,510 symbol images (17.5%), 537 icon images (6.2%), 1,984 ensemble images (23.0%), 971 face images (11.2%), and 1,568 body-part images (18.1%). Because labels were not mutually exclusive, each decoder was fit separately.

#### Decoding

For orthography-related decoding, the MCCA alignment was refit within each outer cross-validation fold using training images only. Held-out images were projected with the fold-specific MCCA weights, and component scores were standardized using training-fold statistics before averaging projections within species. Binary labels were decoded with L2-regularized logistic regression with balanced class weights and 3-fold internal penalty selection, and performance was measured as area under the ROC curve. Cross-validation was grouped by object category, ensuring that exemplars of the same concept did not appear in both training and test sets; the number of folds was set to the largest value up to 5 that maintained valid class balance. Significance was assessed by label permutation, p values were controlled with Benjamini-Hochberg false discovery rate correction across decoders, and species differences were tested with paired fold-wise comparisons.

#### DNN residualization

To test the degree to which orthography-related decoding was explained by low-level visual features, we constructed a control embedding from early layers of the same seven DNNs. For each model, layers in the first 25% of normalized depth were concatenated and, as above, reduced to 250 PCA components within each fold to equalize dimensionality. These early-layer representations were then aligned across models with regularized MCCA using 90 components to match the current universal dimensionality (regularization = 1.0). Within each fold, the species-specific primate predictors were residualized against this embedding using ridge regression with inner cross-validated penalty selection, and decoding was repeated on the residuals.

#### Region-wise analyses

Finally, to assess the progression of orthography-related structure along the ventral pathway, we repeated the alignment and decoding pipeline in V1 and V4/hV4 spaces. Specifically, we refit the same alignment scheme used in IT within each earlier visual ROI, yielding separate cross-species MCCA spaces for V1 and V4/hV4 in addition to IT. These region-specific spaces were built using the same regularization settings and human group-demeaning procedure. We fixed the rank of the V1 and V4 decompositions to the IT-derived rank (k = 90) to avoid cross-region differences being caused by CCA dimensionality. These final decoding analyses were performed on species-averaged projections within each region.

## Supplementary Materials

### Supplementary Note 1: Temporal generalization analysis

To verify that the 75–175 ms post-stimulus window used for macaque IT analyses captures temporally stable category-level structure, we performed a temporal generalization analysis. For each monkey, we trained linear discriminant analysis (LDA) classifiers with Ledoit-Wolf covariance shrinkage to decode 720 object categories from multi-unit activity (MUA) in IT. Classifiers were trained in 20 ms sliding windows (2 ms step) spanning *−*50 to 200 ms relative to stimulus onset and evaluated at all time points in a leave-one-repetition-out cross-validation scheme. The resulting temporal generalization matrices (Fig. S1) confirm robust, temporally sustained category information within the selected analysis window.

**Fig. S1.**
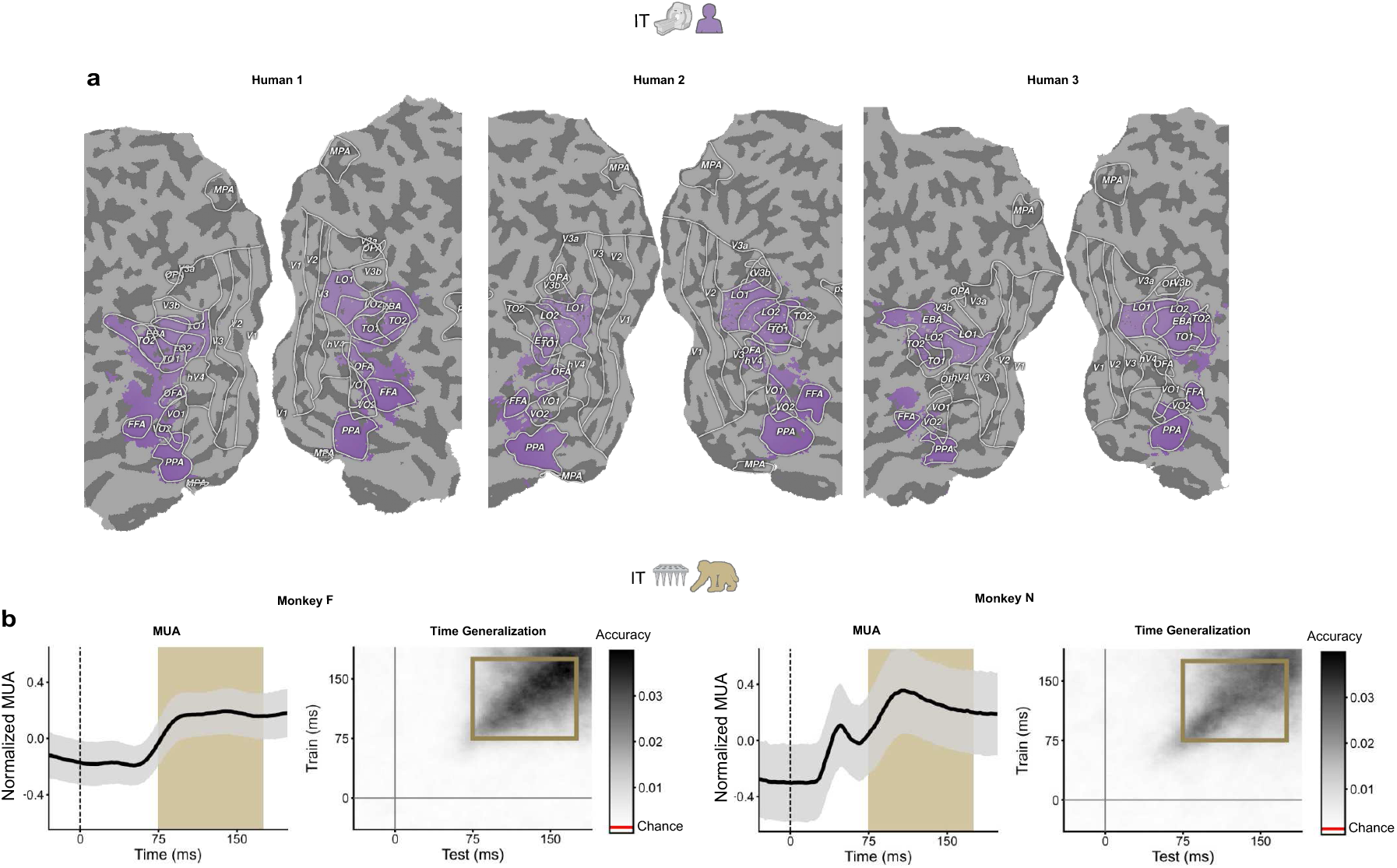
Selected regions and time windows across macaque and human IT. (Top) Cortical flatmaps for each human participant showing the IT region of interest (ROI), defined as the union of LOC, FFA, PPA, EBA, and pRF-defined LO1/LO2 voxels. (Bottom) Mean MUA traces (*±* 1 SD) and temporal generalization matrices for each macaque. The gold shading marks the 75–175 ms analysis window; the square on the generalization matrix marks the corresponding train *×* test region. The red line on the colorbar indicates chance (1/720). See Supplementary Note 1 for details.

**Fig. S2.**
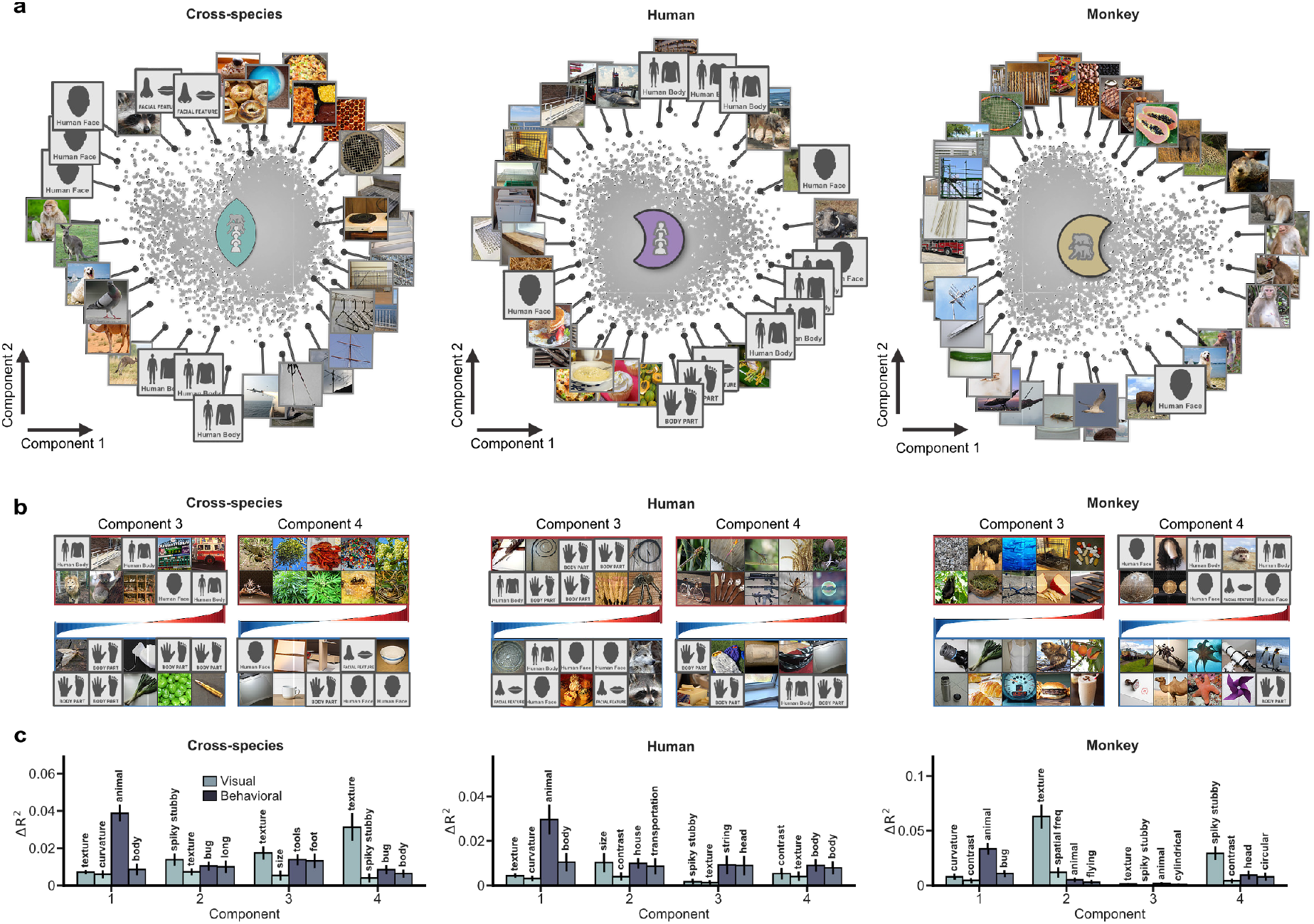
Top image-wise loadings and predictive features of the first four components of each canonical space. (a) Image-wise loadings on the first two components for the cross-species, human-only, and macaque-only MCCA models, with representative high-loading images shown around each scatterplot. (b) Positive and negative pole image grids for components 3 and 4 in each space. (c) Feature-regression results for the first four components show the strongest visual and behavioral predictors, quantified as the drop in cross-validated *R*^2^ when each predictor is removed from the joint model.

**Fig. S3.**
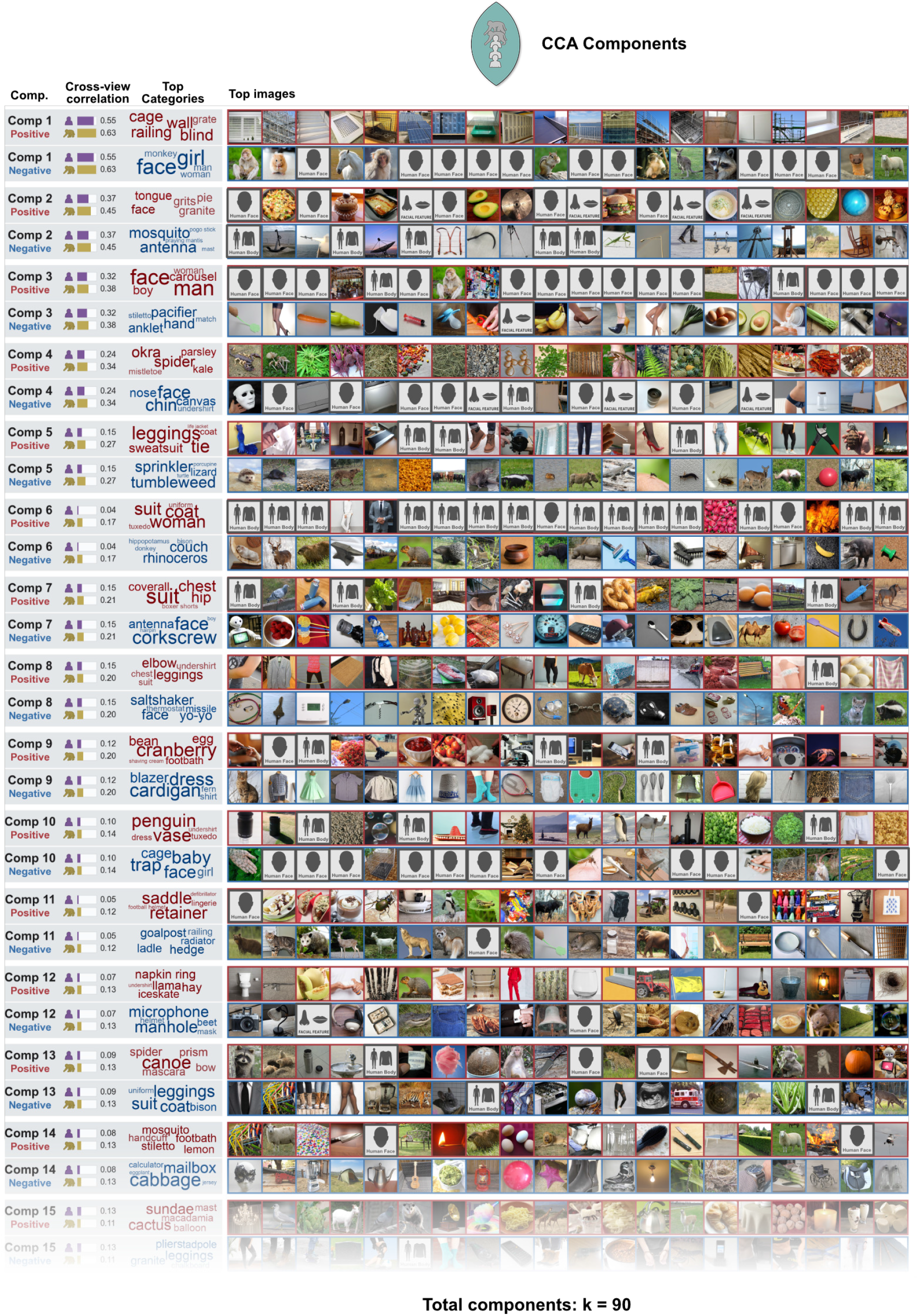
Cross-species CCA component gallery. Gallery of CCA components from the cross-species space. Top components, displayed with their positive and negative poles, held-out cross-view correlation, the four highest-enriched categories, and top-loading images. For image display, images were sampled from the top 1% of stimuli within each pole. The full cross-species space contained 90 retained components.

**Fig. S4.**
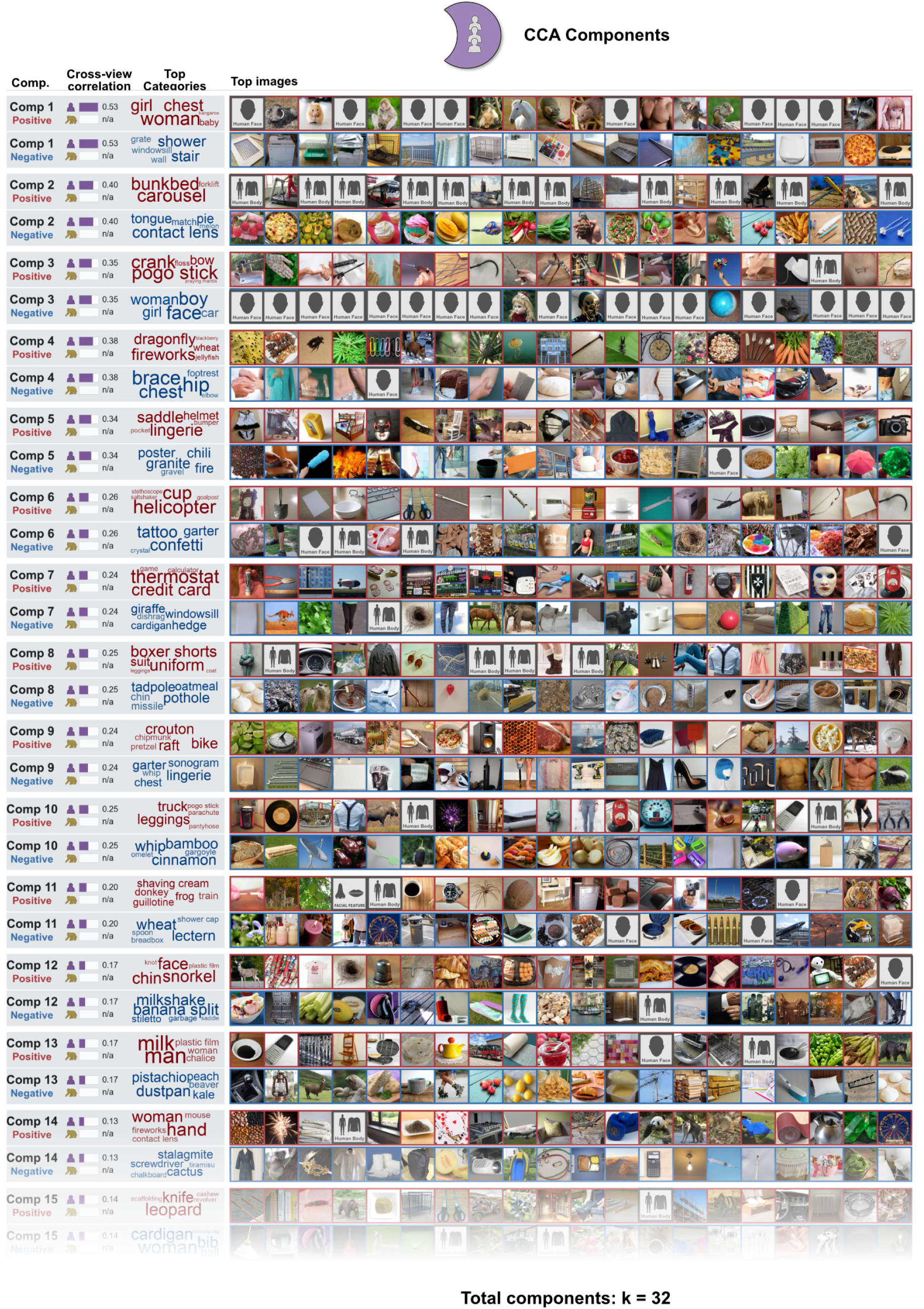
Human CCA component gallery. Gallery of CCA components from the human-only space. Top components, displayed with their positive and negative poles, held-out cross-view correlation, the four highest-enriched categories, and top-loading images. For image display, images were sampled from the top 1% of stimuli within each pole. The human-only space contained 32 retained components.

**Fig. S5.**
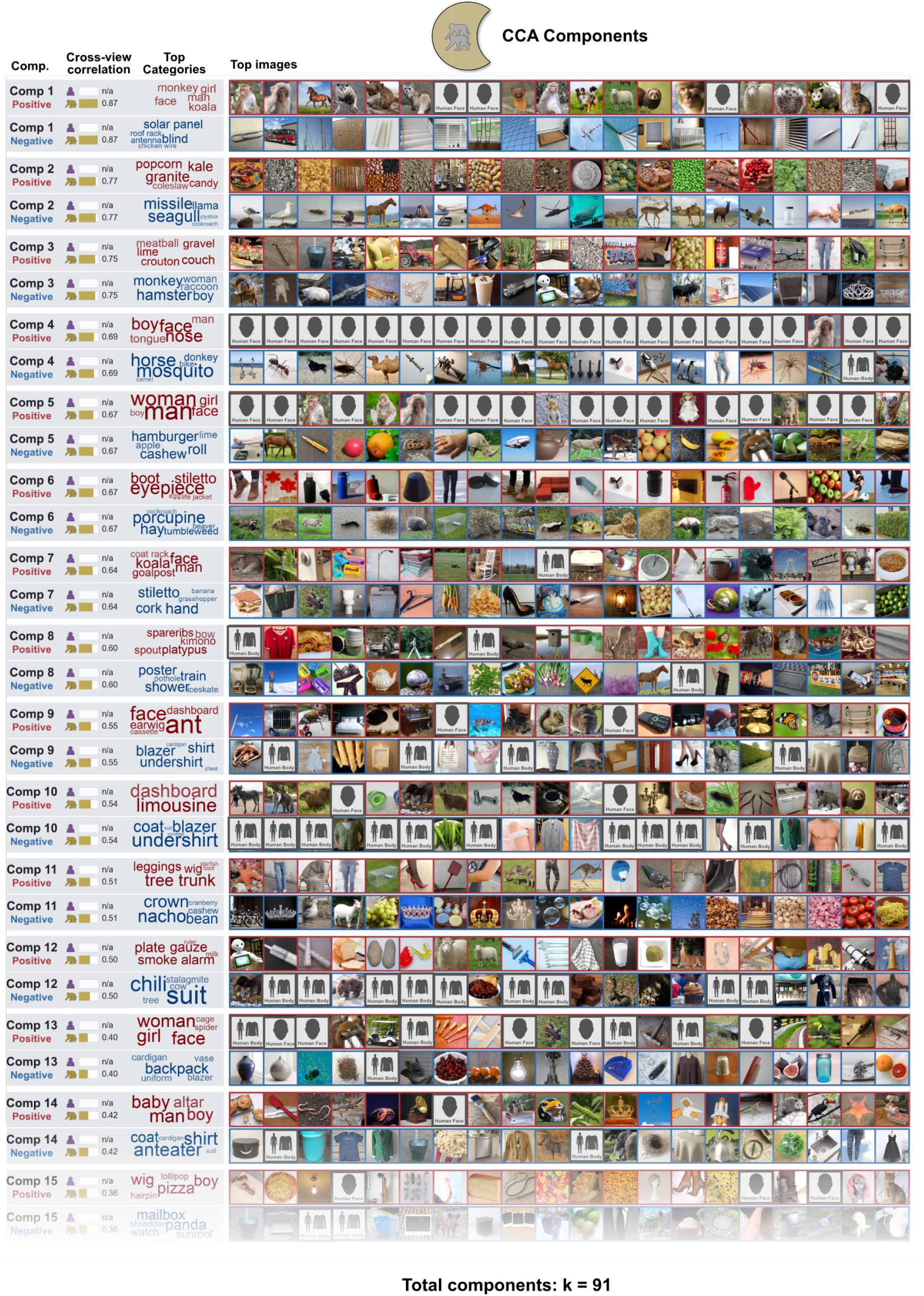
Macaque CCA component gallery. Gallery of CCA components from the macaque-only space. Top components, displayed with their positive and negative poles, held-out cross-view correlation, the four highest-enriched categories, and top-loading images. For image display, images were sampled from the top 1% of stimuli within each pole. The macaque-only space contained 91 retained components.

**Fig. S6.**
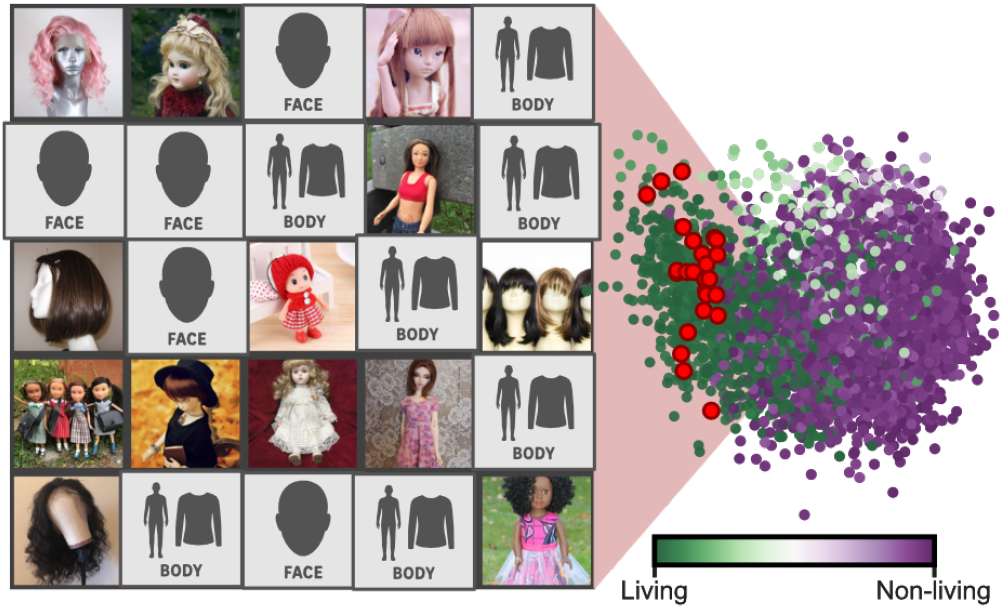
Non-living images loading on the living/animacy pole. (Left) Grid of the 25 non-living images with the most negative loadings on universal canonical component 1 (i.e., the living/animacy pole). These items are semantically non-living but share visual or conceptual properties with animate objects, including puppets, masks, and inanimate objects held or worn by humans. (Right) Scatterplot of all stimulus-wise loadings on universal components 1 and 2, colored by animacy rating. Red circles mark the 25 exception images shown in the left panel.

**Fig. S7.**
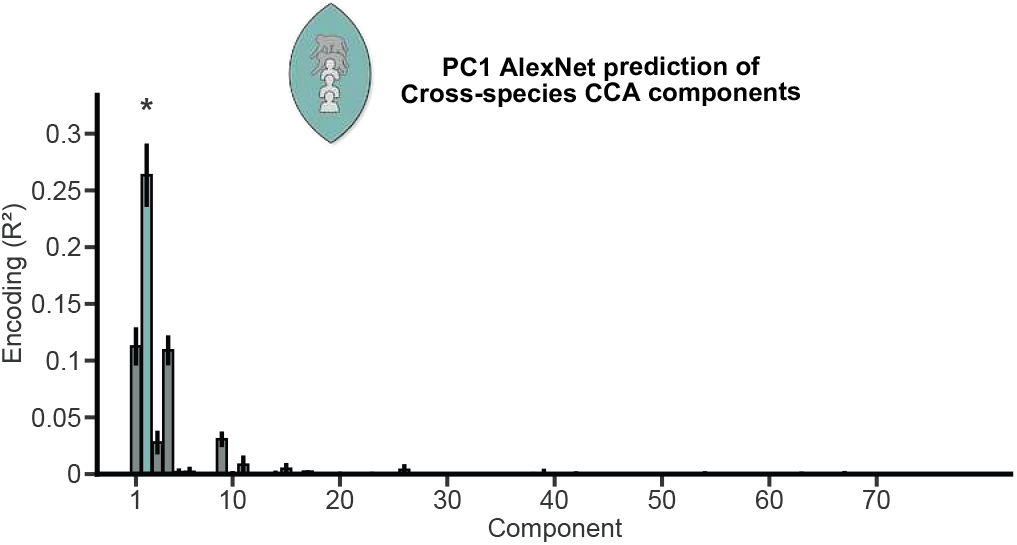
AlexNet fc6 PC1 best predicts the spikiness-related canonical axis. Cross-validated encoding performance (*R*^2^) of AlexNet fc6 PC1 for each universal canonical axis. Component 2 (highlighted) achieves the highest *R*^2^ and significantly outperforms all other components (paired *t*-tests, FDR-adjusted *p <* 0.05), confirming that this axis captures the spikiness dimension previously operationalized via AlexNet fc6 (Bao et al., 2020). Error bars indicate *±* 1 SD across cross-validation folds.

**Fig. S8.**
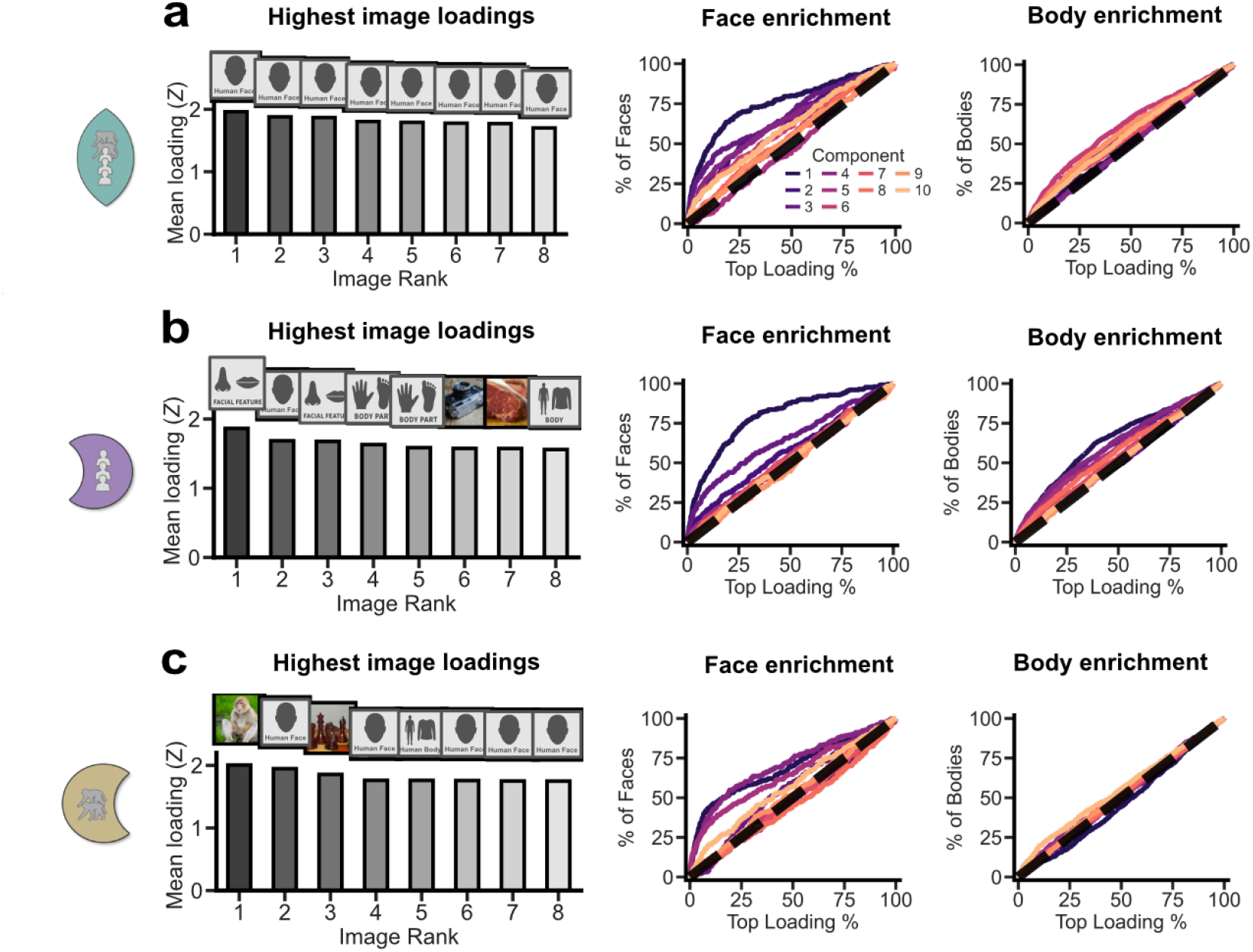
Face and body-part enrichment across leading canonical axes. For the (a) cross-species, (b) human-only, and (c) macaque-only spaces, the left panels show the highest-loading images across the leading canonical axes. The cumulative curves show the fraction of face and body-part images recovered as increasingly large portions of the top-loading stimuli are included for each component. Curves above the diagonal indicate enrichment relative to the full stimulus set.

**Fig. S9.**
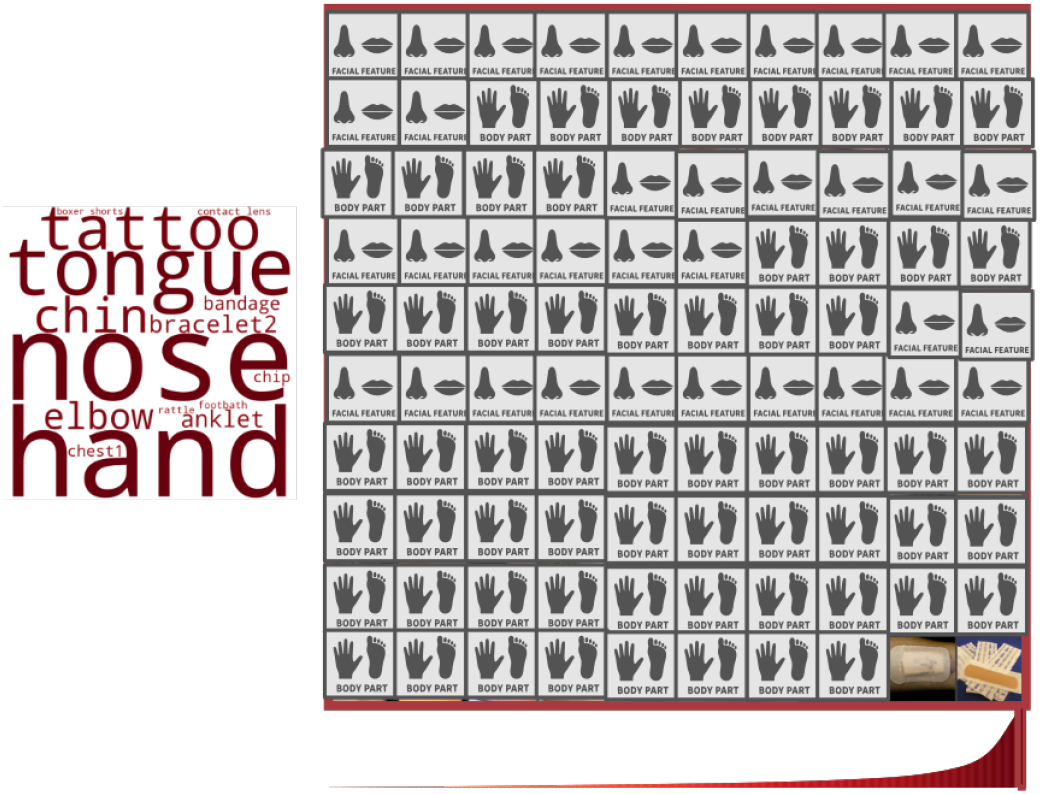
SPoSE body-part component from prior work. Visualization of the behaviorally derived body-part dimension from SPoSE (Hebart et al., 2023). (Left) Word cloud of category-level mean scores on the body-part dimension, with font size proportional to score magnitude. (Right) Mosaic of the 100 highest-scoring images on this dimension, bordered in red. Below the mosaic, a smoothed distribution of all stimulus scores is shown, with the bracket indicating the range of the displayed top images.

**Fig. S10.**
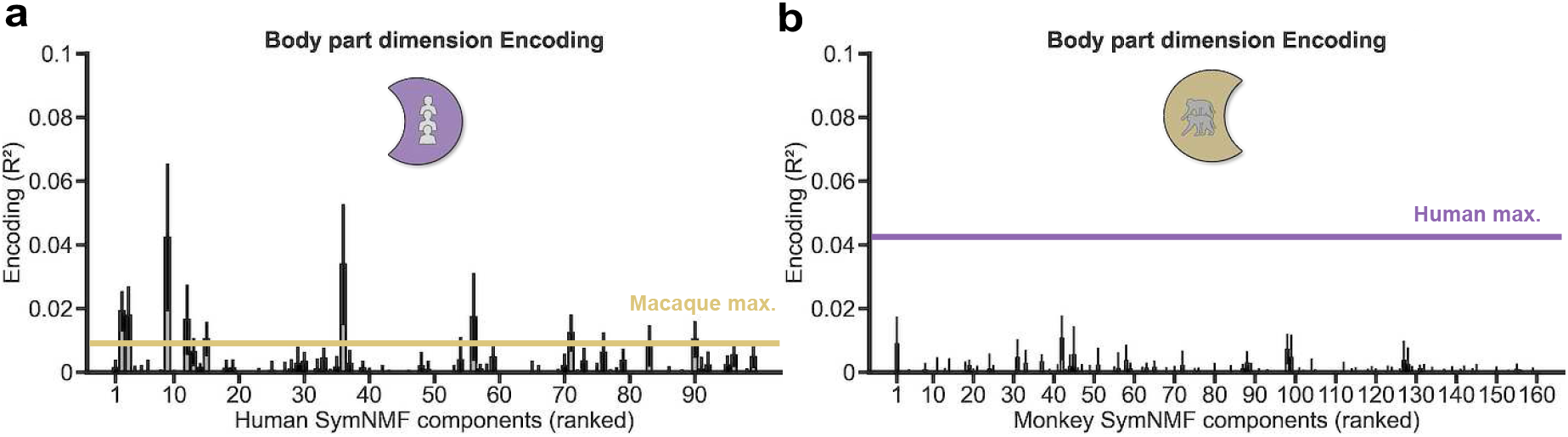
Body-part covariate analyses for species-specific sNMF factors. Cross-validated *R*^2^ for predicting each species-specific sNMF factor from the behavioral body-part dimension (Hebart et al., 2023) using ridge regression. (a) Human-selective factors, ranked by *R*^2^. (b) Macaque-selective factors ranked by *R*^2^. Error bars indicate *±* 1 SD across folds. Horizontal lines reflect maximum body part encoding values of the other within-species space.

**Fig. S11.**
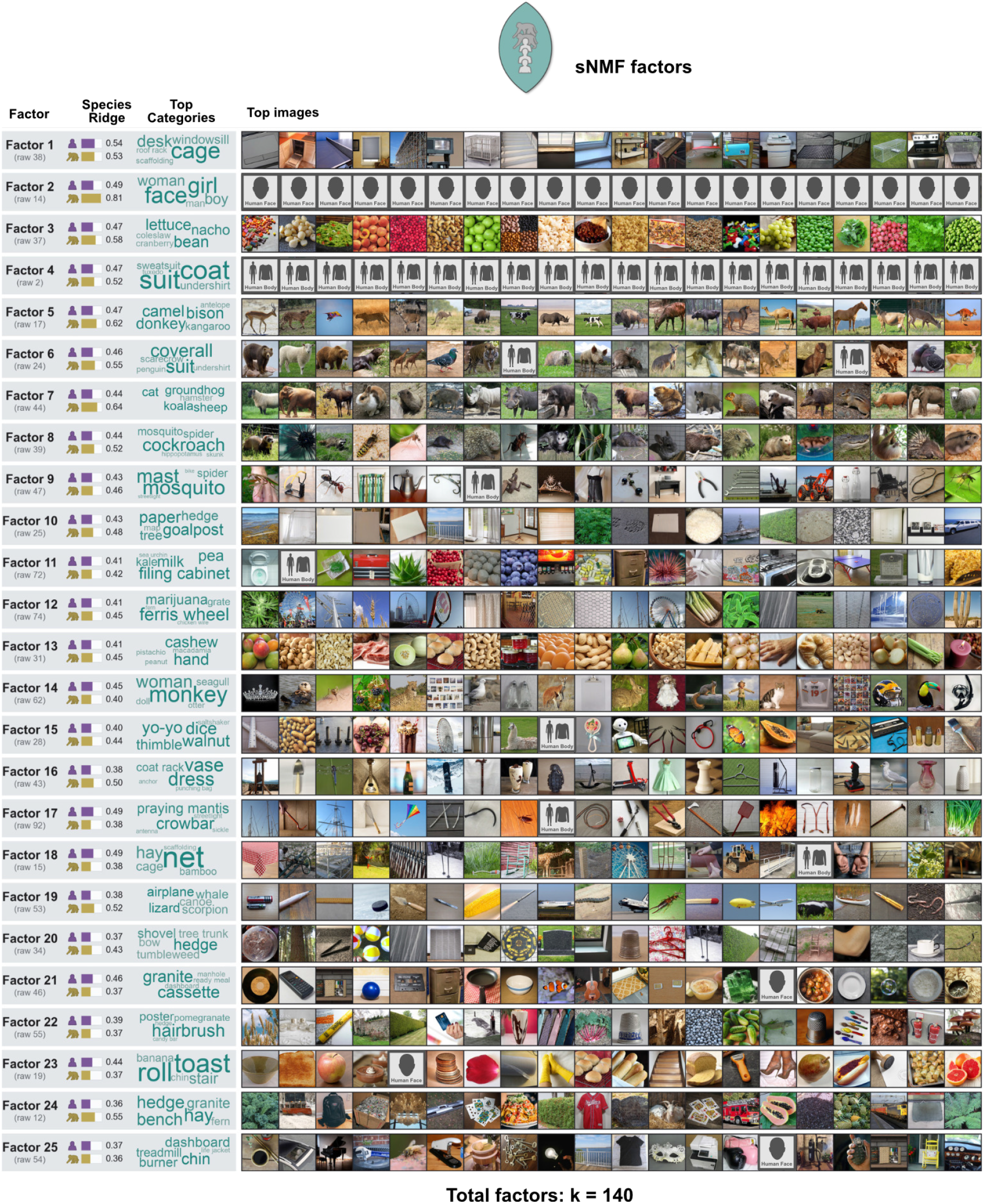
Shared sNMF factor gallery. Gallery of sNMF factors from the universal space. Each row shows one factor with its rank, cross-validated *R*^2^ per species, top enriched categories, and top 20 images. For readability, only the top 25 ranked factors are shown from the full retained factor set. Factors were retained if they were recoverable from the human and macaque projections onto the cross-species CCA space (*R*^2^ *≥* 0.20), and are ranked by minimum *R*^2^ across the two species.

**Fig. S12.**
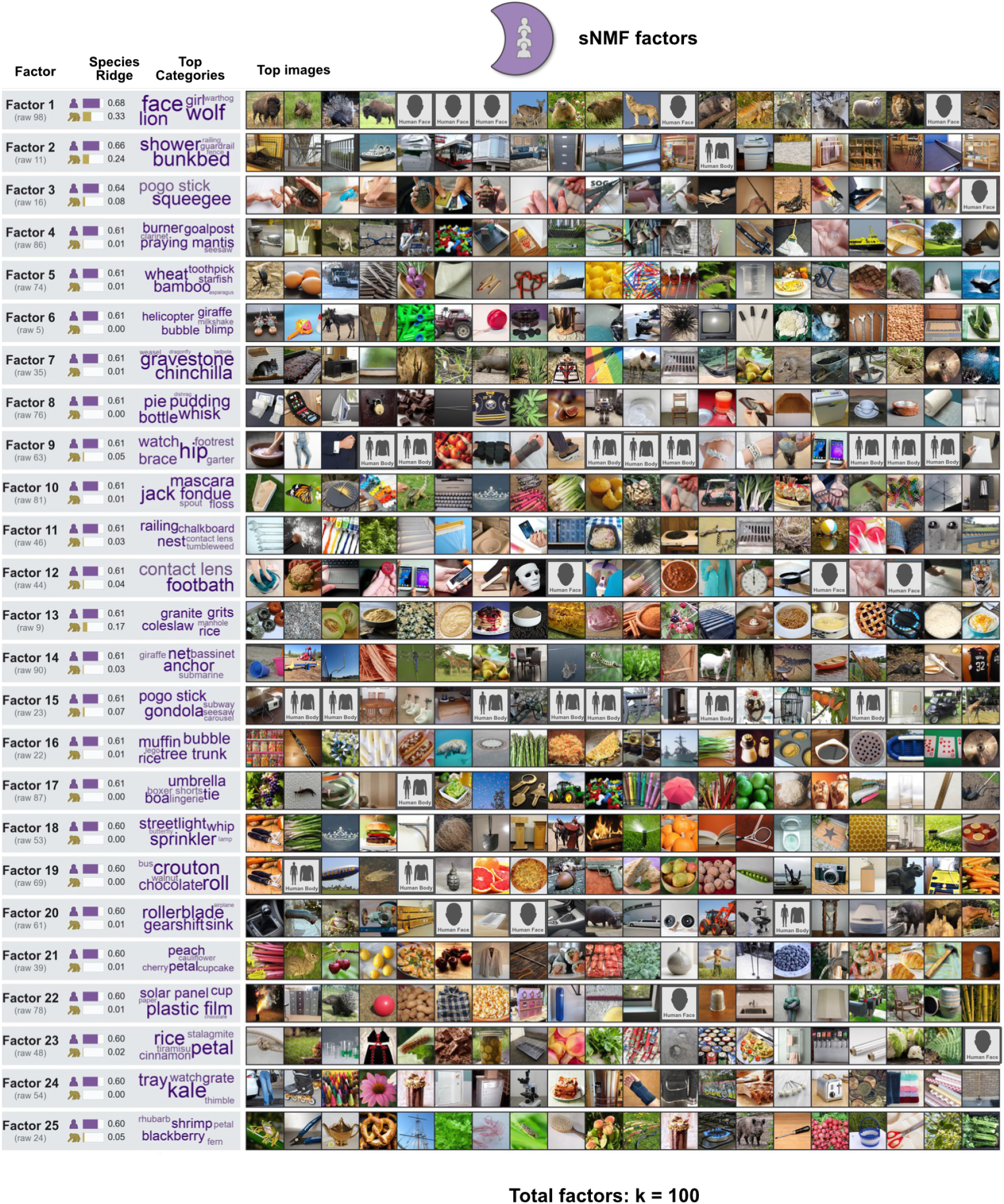
Human sNMF gallery (human-selective factors). Gallery of human-selective sNMF factors from the within-species decomposition. Each row shows one factor with its rank, cross-validated *R*^2^ per species, top enriched categories, and top 20 images (same layout as Fig. S11). For readability, only the top 25 ranked factors are shown from the full retained factor set. Factors were retained if they were recoverable from the human CCA space (*R*^2^ *≥* 0.20) and substantially weaker in the macaque space (macaque *R*^2^ *<* 50% of human *R*^2^), and are ranked by human *R*^2^.

**Fig. S13.**
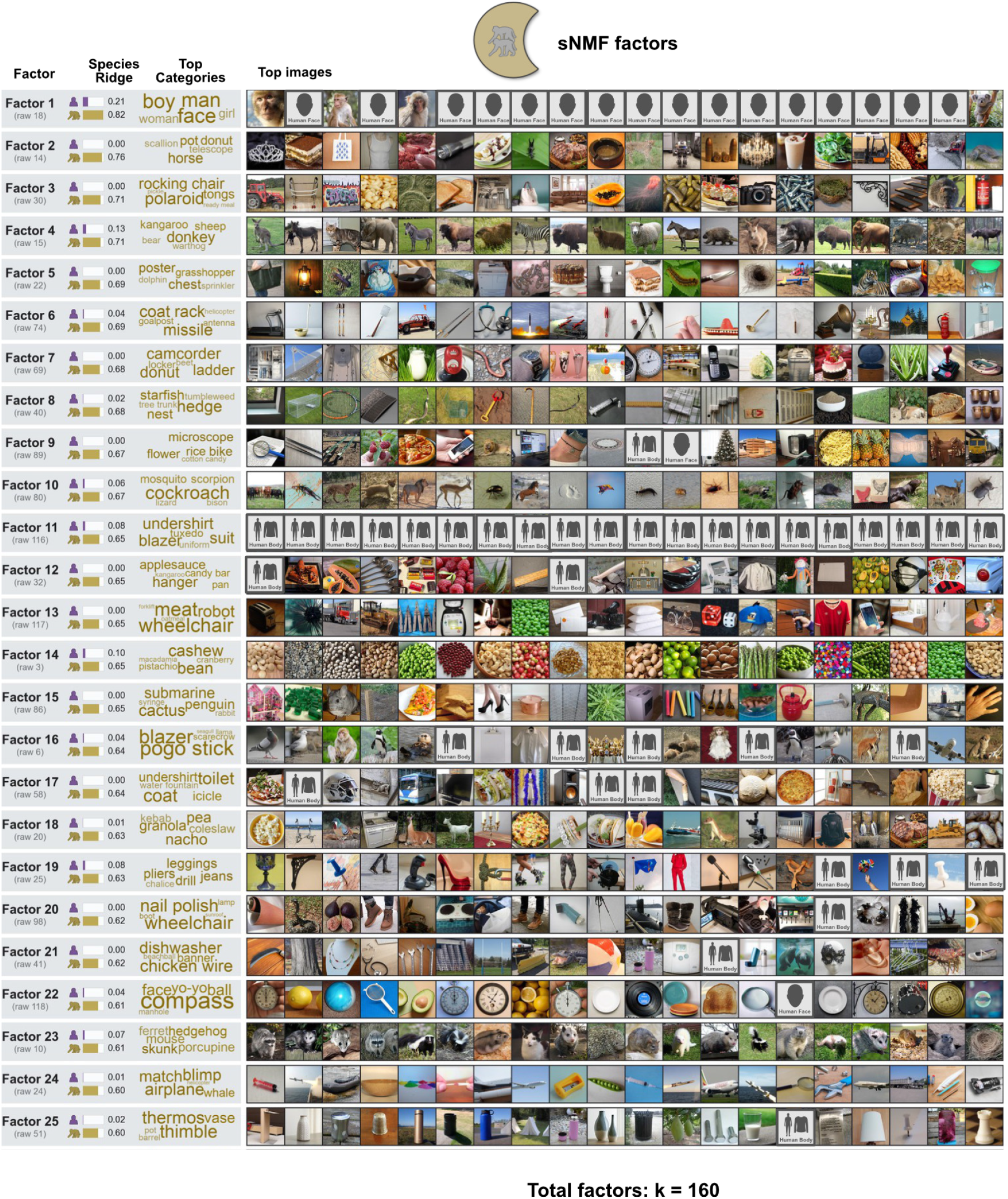
Macaque sNMF gallery (macaque-selective factors). Gallery of macaque-selective sNMF factors from the within-species decomposition. Each row shows one factor with its rank, cross-validated *R*^2^ per species, top enriched categories, and top 20 images (same layout as Fig. S11). For readability, only the top 25 ranked factors are shown from the full retained factor set. Factors were retained if they were recoverable from the macaque CCA space (*R*^2^ *≥* 0.20) and substantially weaker in the human space (human *R*^2^ *<* 50% of macaque *R*^2^), and are ranked by macaque *R*^2^.

**Fig. S14.**
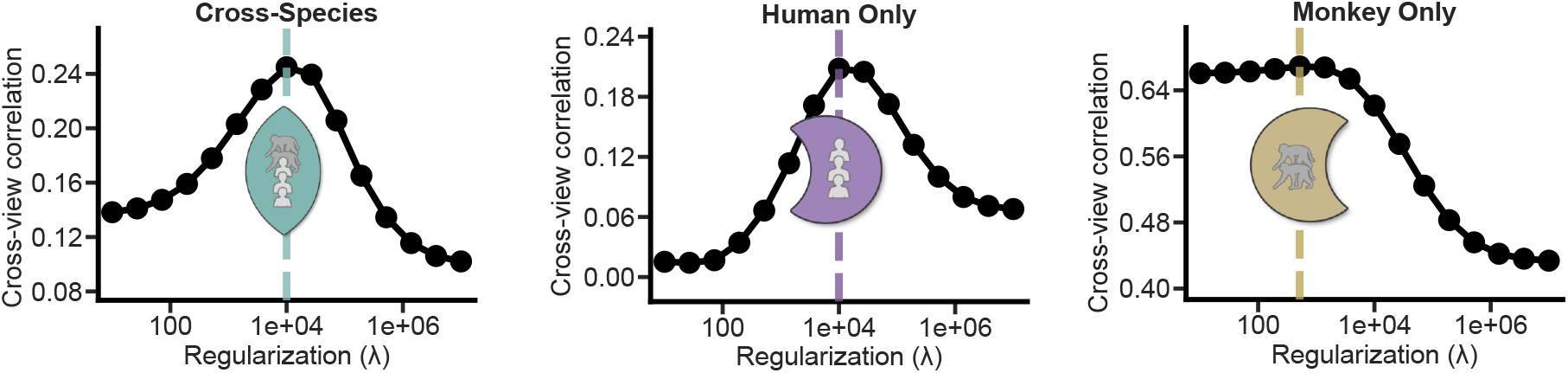
MCCA regularization-selection results across model families. Regularization parameter selection for each MCCA model family (cross-species, human-only, macaque-only). Each panel shows cross-view similarity (Pearson’s *r*) as a function of the ridge regularization parameter *λ* across a log-spaced grid (*λ ∈* [10^1^, 10^7^]). The dashed vertical line indicates the selected *λ* value, chosen to maximize mean held-out cross-view correlation in nested cross-validation.

